# Single-cell Genomic Copy Number Evolution Reveals Frequent Loss of the Y chromosome in Esophageal Adenocarcinoma

**DOI:** 10.1101/2025.09.14.676175

**Authors:** Shiwei Yin, Na Zhong, Swati Agrawal, Bingru Feng, Sneha Pradhan, Nathan K. Jobalia, Maryam Vaziripour, Sampath K. Poreddy, Laura A. Kresty, Erin A. Gillaspie, Nicholas E. Dietz, Waddah B. Al-Refaie, Kalyana C. Nandipati, Yusi Fu, Jun Xia

## Abstract

The incidence of esophageal adenocarcinoma (EAC), an aggressive cancer, is rising in Western countries, and incidence is higher in men. To better understand the roles of DNA copy number evolution, tumor heterogeneity, and tumor microenvironment in EAC and its origins, we evaluated EAC, Barrett’s Esophagus (BE), and normal adjacent tissue using our previously developed ultra-sensitive, single-nucleus DNA copy number analysis (12,186 nuclei, 20 specimens) and single-nucleus RNA-sequencing (snRNA-seq; 87,027 nuclei, 32 specimens). Loss of Y (LOY) chromosome was rare in BE but markedly enriched in EAC. In genome unstable cells, complete LOY chromosome often coincided with X-chromosome duplication. We did not detect X-chromosome inactivation via *XIST* expression in cells with LOY, suggesting increased expression of X-chromosome genes and potential effects on tumor immune evasion. When considering patients’ clinical features, we identified distinct differences in tumor microenvironment by sex, age, and obesity status, particularly massively reduced immune cell composition in obese patients. Finally, our cell line models recapitulate LOY, and we identify the first candidate anti-LOY agent. Our findings highlight how LOY affects tumor subpopulations and shapes the tumor microenvironment during EAC progression.

## Main Text

Esophageal adenocarcinoma (EAC) is a highly aggressive malignancy characterized by low overall 5-year survival rate of about 20% (*1*). EAC incidence has surged by 700% from 1973 to 2014 (*2*) and is much higher among males than females(*3*). This disparity between sexes is poorly understood (*4*). Bulk sequencing has identified many common genetic alterations in EAC, such as *TP53* mutations and *c-Myc* amplification (*5-9*). One substantial genetic alteration, loss of the Y chromosome (LOY), has been observed in cancer (*10*), aging(*11*), and cardiomyopathy(*12*), but its causes, mechanisms, and consequences remain unclear (*13*). Existing methods for detecting LOY have limited accuracy and sensitivity (*14*), due in part to tumor heterogeneity. Thus, the landscape and functional impact of LOY in clinical specimens have not been fully defined, particularly for diseases with a higher incidence in men, like EAC. Advancements in single-cell genomics have provided deeper insights into tumor biology by enabling the study of cellular heterogeneity, but the complexities of inter- and intra-tumor heterogeneity at DNA copy number (CN) and transcriptional levels remain underexplored (*7*).

Cancer progression is also influenced by the tumor microenvironment (TME), which can significantly impact tumor growth, metastasis, and therapy response (*15*). The TME includes many cellular types and components (e.g., immune cells, fibroblasts, endothelial cells, extracellular matrix, and signaling molecules). Single-cell sequencing lets us examine the effects of both CN alterations within tumors and the TME by interrogating the gene expression profiles of different cellular components (*16*). To date, the impact of CN alterations and their interactions with the TME in EAC remain largely unknown (*17, 18*). Thus, there is an urgent need to explore the role of CN alterations, particularly LOY, and TME in driving EAC progression.

In this study we used an ultra-sensitive, single-cell, DNA CN analysis we developed previously, called <u>Ultra</u>-sensitive single cell/nucleus <u>C</u>opy <u>N</u>umber <u>A</u>lteration sequencing (Ultra-CNA) (*18*), and snRNA-Seq to examine EAC tumors and paired tissues. We identified critical events, including LOY in tumor and adjacent tissues, that potentially drive early EAC progression. By integrating genomic and transcriptomic data at the single-cell level, we identified interplay between tumor cells and the TME, revealing how LOY drives genome instability and dysregulates the immune system. This information could facilitate the selection of and predict response to treatment.

### Single-cell Copy Number Landscapes of Esophageal Adenocarcinoma

Genomic CN alterations are significant predictors of the transition from Barrett’s Esophagus (BE) to EAC (*6, 7*). CN patterns can potentially identify patients at risk of developing cancer years before traditional markers, like dysplasia. CN alterations are often masked from detection because of intra-tumor heterogeneity and methodological limitations(*19*). Thus, we need more sensitive approaches to detect CN alterations in a heterogeneous tumor background and to inform the development of new biomarkers and elucidate the mechanisms implicated (*20*). We leveraged Ultra-CNA (*18*), which detects CN alterations as small as 10kb (Figs. 1A, B), to profile 12,186 single nuclei from 20 specimens across 11 patients, including samples of EAC, BE, and normal squamous mucosa located at least 5 cm from a neoplasm site (Fig. 1C). This comprehensive dataset enabled the capture of significant genomic alterations within and between samples at single-cell resolution. After applying additional filtering criteria (mapping rate >50%, >100k reads), we generated heatmaps displaying CN alterations for each specimen (Figs. 1D and S1–11). The heatmaps revealed extensive CN variability between cells in the same sample, verifying significant tumor heterogeneity. Specifically, EAC samples showed markedly higher genomic instability than BE samples (Figs. 1D, 2, and S1–11), consistent with previous findings that linked elevated genomic alterations to aggressive EAC progression (*21*).

**Fig. 1.**
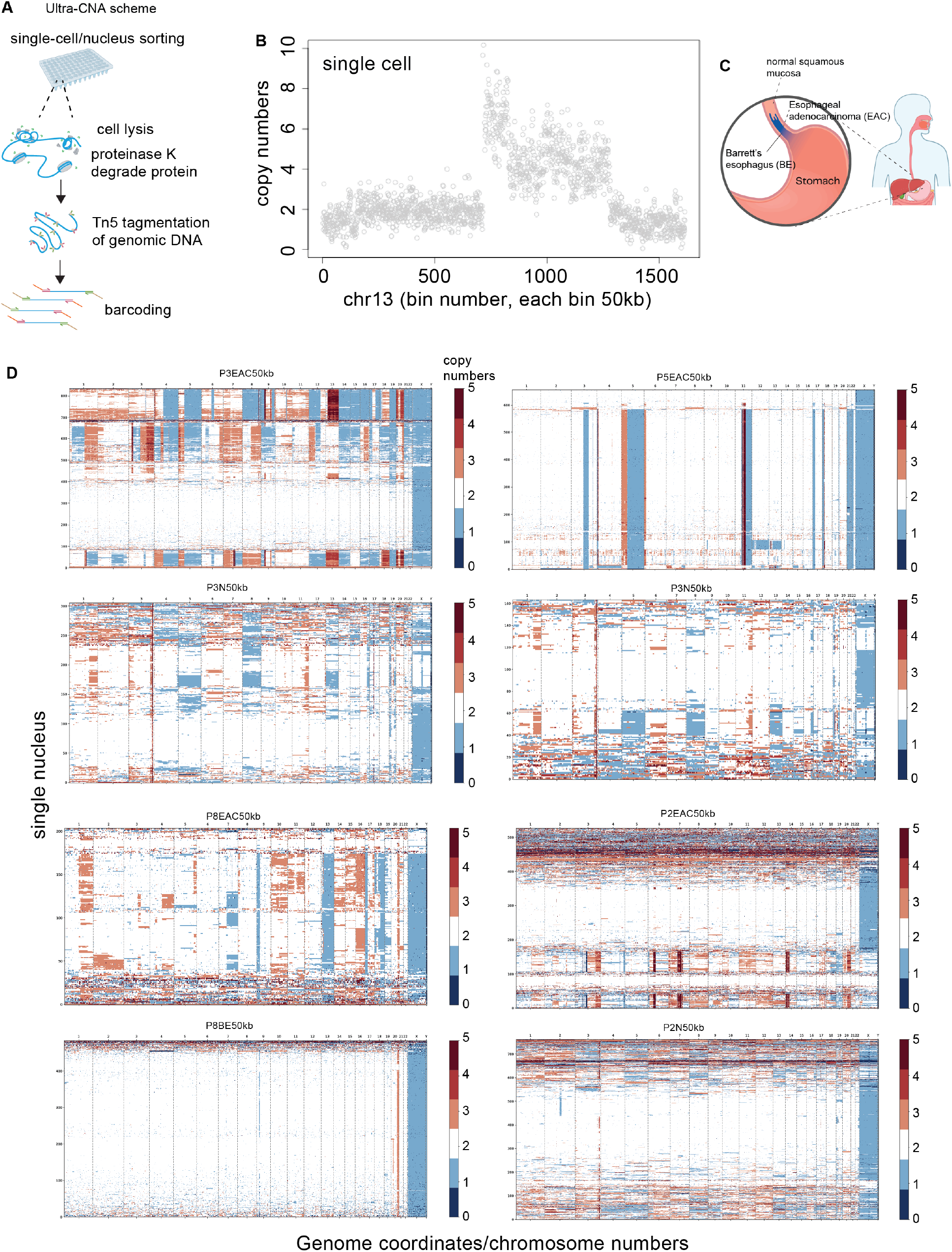
Single-cell Copy Number Landscapes of Esophageal Adenocarcinoma (EAC). (**A**) Overview of the Ultra-CNA workflow to detect single-cell/nucleus copy number alterations with high sensitivity. (B) A demonstration of Ultra-CNA’s capability to identify copy number gains in individual 293T cells with high resolution. (C) Collection sites for specimens representing EAC, Barrett’s Esophagus (BE), and adjacent normal squamous mucosa. (D) Single-nucleus copy number profiles for representative EAC, BE, and normal specimens, illustrating the genomic complexity and instability within each specimen.

**Fig. 2.**
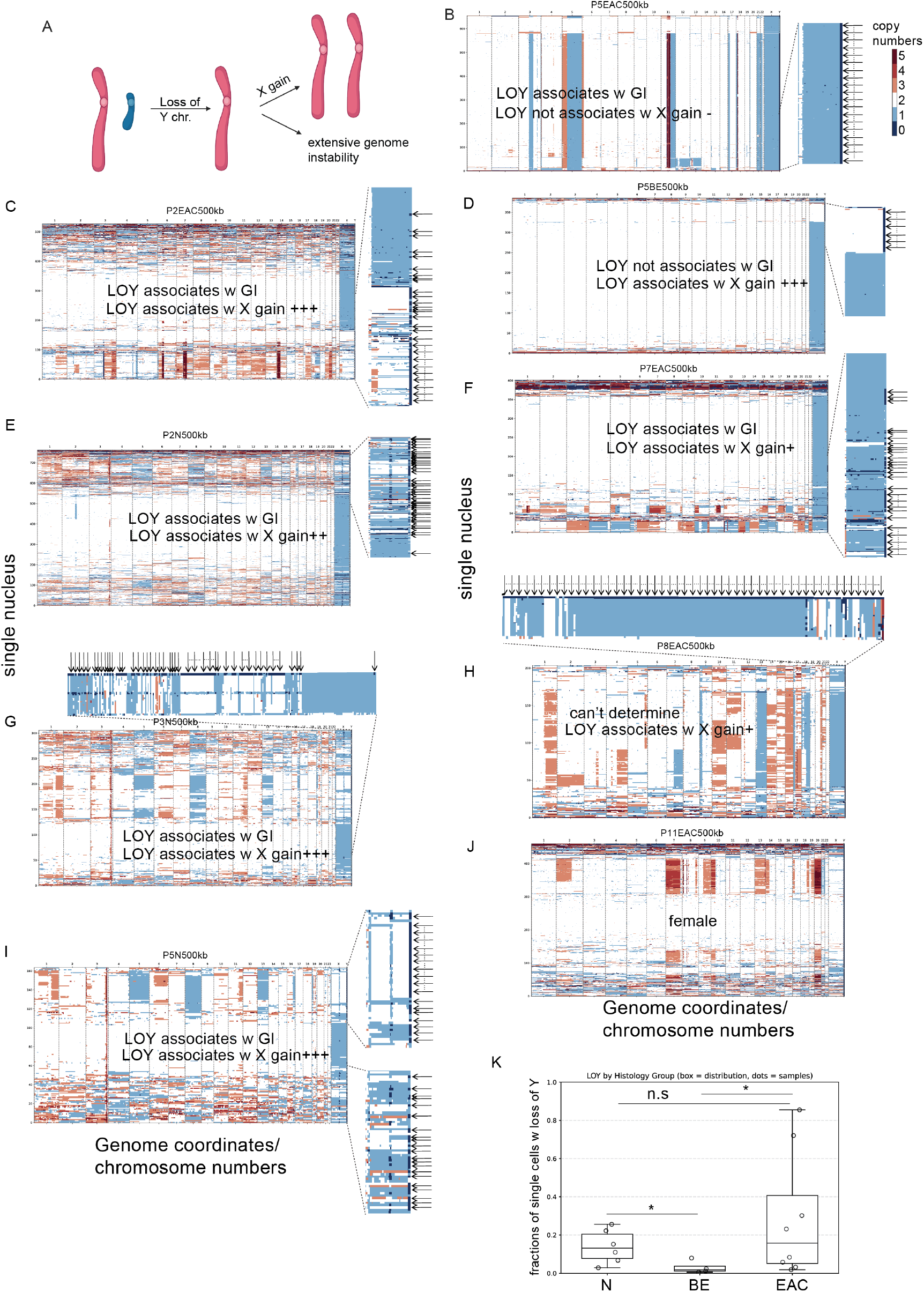
LOY Association with X-Chromosome Duplication and Genomic Instability. (**A**) Model illustrating the frequent loss of Y chromosome (LOY) due to an unknown mechanism, often accompanied by X-chromosome duplication and associated with extensive genomic instability. (B–I) Genomic profiles of EAC, BE, and normal squamous mucosa specimens reveal frequent LOY. In 75% (6/8) of cases, LOY is associated with genomic instability, and in 87.5% of specimens (7/8), LOY is linked to gain of X chromosomes. (J) Specimen from a female EAC patient showing cells with predominantly two X chromosomes and no Y chromosomes. (K). LOY is more frequent in EAC than BE. Normal squamous mucosa from cancer patient has higher LOY than BE. Turkey boxplots. Mann Whitney U test.

### LOY in EAC and Adjacent Normal Squamous Mucosa

In our analysis of male patients diagnosed with EAC, we frequently observed LOY in a subpopulation of cells. Specifically, out of 8 male EAC patients with Ultra-CNA data, 4 tumors (50%) had with >20% of cells showing complete LOY (Figs. 2 and S1–11). This finding suggests that LOY is a frequent and potentially critical genomic alteration in EAC. Frequent LOY is seen in other cancers, such as bladder and lung cancer (*22*), supporting the hypothesis that LOY can be a common event in male cancer patients, particularly in cancers originating from epithelial tissues and those attributed to environmental exposures. In addition to complete LOY, our analysis revealed frequent X-chromosome duplications (Figs. 2A, model, and 2B–J). There were relatively few instances of partial LOY. Unexpectedly, these changes also appeared in adjacent squamous mucosa-derived cells (Fig. 2K) confirmed to be normal squamous esophageal mucosa without any features of cancer (Fig. S12, P2N) based on review of hematoxylin & eosin-stained sections. These results suggest that LOY can arise as a very early, preneoplastic somatic event in histologically normal esophageal epithelium. While we cannot determine whether these LOY events were induced by treatment with chemo- and/or radiotherapy or if it arose spontaneously, the observation that only a subset of BE and normal squamous mucosa specimens from treated patients exhibit LOY argues in favor of a spontaneous origin.

Finding LOY beyond the tumor compartment implies that broader genomic instability in the esophageal epithelium could precede or contribute to oncogenesis. Although LOY has been reported in various cancers, the discovery of complete LOY within specific subpopulations together with accompanying X-chromosome duplications has not been described previously both in and outside of cancer tissue. To corroborate these findings, we used fluorescence in situ hybridization (FISH) to detect LOY and other CN alterations on chromosomes X, Y, and 1 in addition to checking the expression of genes on sex chromosomes (Fig. 3). We used validated probes targeting DXZ, DYZ1, and the CEN1p13.3 region (methods), marking the X and Y chromosomes and chromosome 1 centromere, respectively. FISH analyses revealed LOY in both EAC and histologically normal squamous mucosa (Figs. 3A, S12-13), confirming that LOY is not confined to malignant cells. Additionally, the frequency of single cells with LOY was significantly higher in EAC samples than BE samples [mean 28.8% vs 3.0%, Figs. 2K, 3A and S1–11, *p*=0.0485, Mann Whitney U test], indicating cells with LOY are enriched during malignant transformation.

**Fig. 3.**
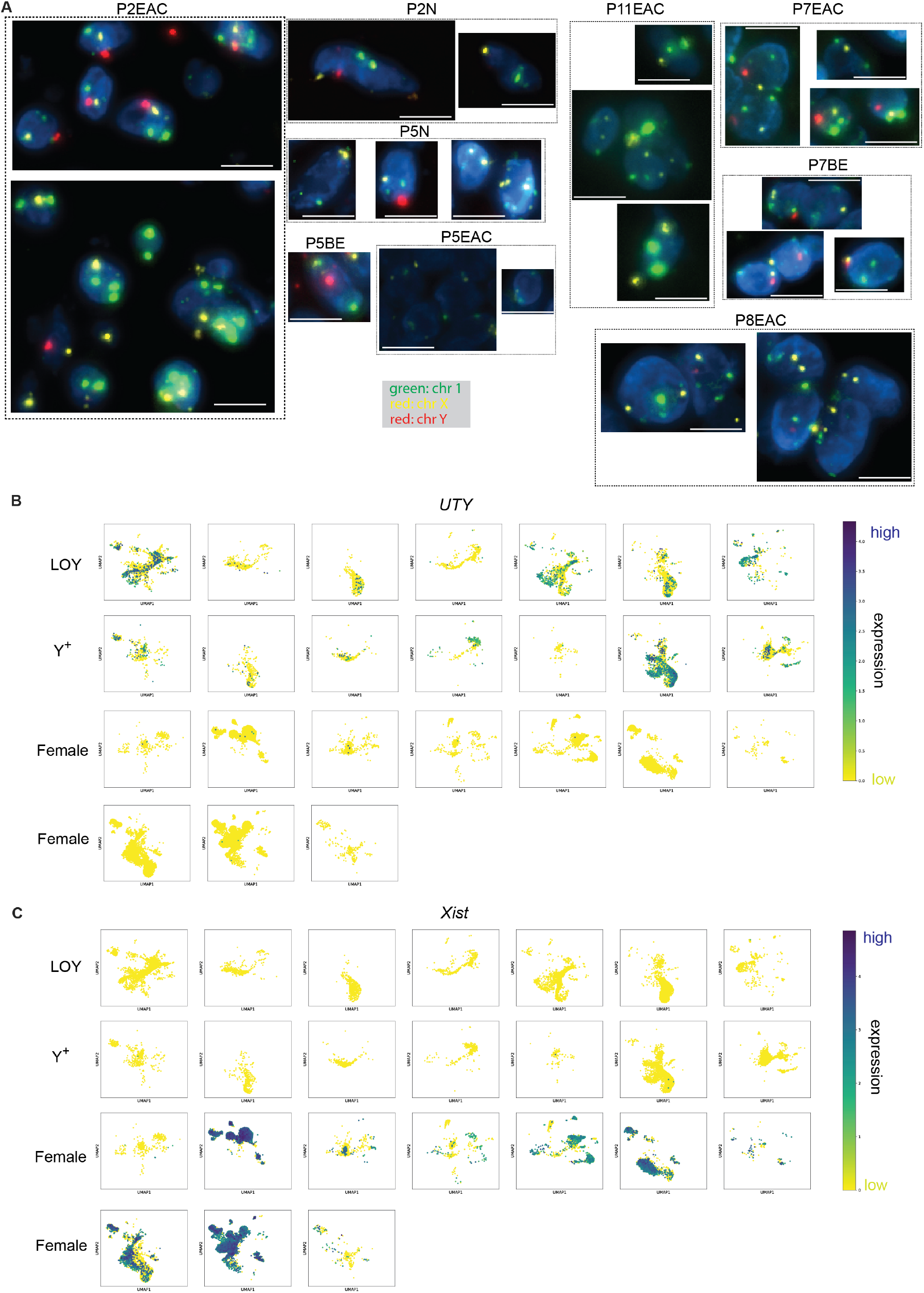
Confirmation of LOY and X-chromosome Duplication in EAC/BE Specimens. (**A**) fluorescence in situ hybridization analysis validates LOY and X-chromosome duplication in samples identified by single-cell Ultra-CNA. Green: chromosome 1, yellow: chromosome X, red: chromosome Y. (B–C) Single-cell RNA-seq analysis of *UTY* and *XIST* expression in male-derived samples with LOY (LOY) or intact Y chromosome (Y+) and female-derived samples. Darker colors indicate higher expression levels. Each box: individual specimen, each dot: cell.

### LOY is Associated with X-Chromosome Duplication and a Lack of *XIST* Gene Expression

Cells with LOY and X-chromosome duplication presented a gene dosage landscape fundamentally different from those of canonical X-chromosome duplication states, such as De la Chapelle syndrome, where one X chromosome is transcriptionally silenced via *XIST*-driven X-chromosome inactivation(*23*). FISH and CN profiling of our specimens found a subset of XX male cells among those with LOY alone (Fig. 3A), yet RNA-seq showed virtually no *XIST* expression (Fig. 3C), indicating that both X chromosomes are transcriptionally active. The Y-chromosome–specific *UTY* gene was expressed in most male-derived cells (Fig. 3B), including those with XX genotypes discovered somatically. This expression is presumably from the subset of cells with the Y chromosome intact. The observed LOY did not arise from the germline, given that shallow whole-genome sequencing of peripheral blood mononuclear cells from our study cohorts found normal Y-chromosome dosage in male patients (Fig. S14). To confirm that contamination was not the cause of the apparent X-chromosome duplication in LOY cells, we used aggregated single-nucleotide polymorphism analysis to ensure the cells came from their associated EAC patient, regardless of whether there was one or two X chromosomes (Fig. S15).

This unique configuration could offer a selective advantage for cancer cells: (i) it compensates for the loss of essential Y-linked functions by increasing expression of corresponding X-linked paralogues and (ii) it simultaneously increases the CN of X-encoded cancer-associated genes [e.g., *TMSB4X*(*24*)], potentially accelerating malignant transformation. Compared with De la Chapelle syndrome (*23*), where X-chromosome inactivation restores near-physiological X-gene dosage, our data point toward a gain in X-gene dosage, which could be an early, cell-intrinsic driver of EAC development. Thus, these X duplicated cells could indicate elevated risk of EAC development. Further, these observations provide mechanistic insights into how aneuploidy-induced dosage imbalances could initiate or propel tumorigenesis.

### Single-nucleus Transcriptomic Landscape of EAC

In addition to evaluating CN changes, we performed snRNA-seq on nuclei from a subset of paired samples to characterize the cellular composition and expression profiles of individual cell subclusters. In total, we analyzed 87,027 nuclei from 32 specimens encompassing EAC, BE, and normal esophageal tissues (with a subset matched, Table S1). When visualized with uniform manifold approximation and projection clustering (Figs. 4A–C), we found distinct populations of epithelial cells, T cells, B cells, and other cell types, underscoring significant transcriptomic heterogeneity within each sample (Fig. 4C).

**Fig. 4.**
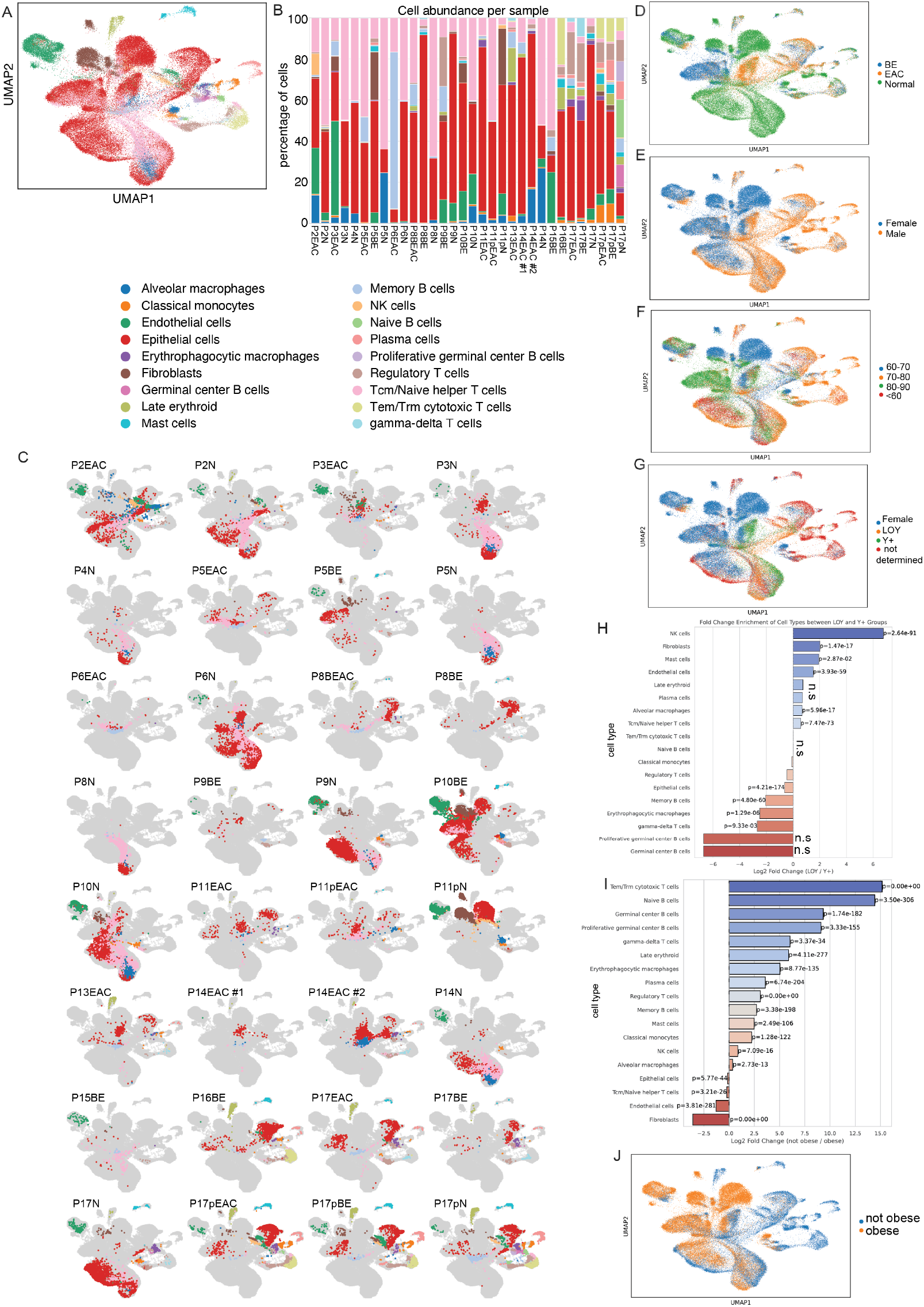
EAC Single-cell RNA-sequencing (scRNA-seq) Landscape. An integrated analysis of scRNA-seq data across various conditions. (**A**) Uniform manifold approximation and projection (UMAP) clustering reveals the distribution of major cell types, including epithelial cells, T cells, and B cells, based on transcriptomic profiles. (B) Stacked bar plot illustrating the proportions of cell types in each patient specimen. (C) The per-sample UMAP plot shows the distribution by cell type. (D) The UMAP plot highlights the separation of cells derived from BE, EAC, and normal tissues. (E) Sex-based distribution, showing male (blue) and female (orange) cells. (F) Age-based distribution across various age groups. (G) UMAP plots comparing Y-chromosome status, identifying LOY (red), intact Y chromosome (Y+, blue), and female (orange) groups. (H) Fold-change differences in gene expression between LOY and Y+ cells. Fisher’s exact test with BH adjustment. (I) Fold-change differences in gene expression between individuals who are and are not obese. Fisher’s exact test with BH adjustment. (J) A UMAP plot comparing the not obese (blue) and obese (orange) groups.

We observed marked differences in immune cell composition across BE, EAC, and normal tissues (Figs. 4D and S16). Notably, γδ, effector memory, and tissue-resident memory T cells were increased in both EAC and BE (Fig. S16), whereas naïve, proliferative germinal center, and germinal center B cells were markedly reduced (Fig. S16). NK cells were increased in EAC samples and reduced in BE samples compared with normal mucosa (Fig. S16), suggesting antitumor response is potentially stronger in EAC. The BE reduction is consistent with prior reports (*25*), whereas prior evidence regarding NK-cell abundance in EAC remains inconclusive(*26*). This immune infiltration and the intricate interactions among different immune cell subsets reflect state-specific changes during carcinogenesis.

Next, we investigated how sex and age impact the TME. Analysis by sex revealed that transcriptomic profiles were distinct between samples from male and female participants (Fig. 4E). Specimens from female participants had higher proportions of fibroblasts and endothelial cells and depleted B cells compared with those from male participants (Fig. S17), indicating possible differences in immune and stromal composition based on sex. Additionally, analysis by age showed increased γδ T cells in older age groups (Figs. 4F and S18), highlighting how aging could affect the TME in EAC(*27*).

Next, we investigated the influence of LOY on immune cell composition. Samples with LOY (defined in Table S2) were significantly enriched in NK cells (p=2.64×10^−91^) and depleted in various B-cell subsets. We observed smaller reductions in γδ T cells and erythrophagocytic macrophages (Figs. 4G–H). These observations are consistent with previous reports linking LOY to altered immune profiles and immune escape(*10*). Notably, we did not detect an increase in regulatory T cells or upregulation of T-cell exhaustion markers(*10*) (Fig. S19). This finding differs from those of mouse bladder cancer models suggesting that LOY drives T-cell exhaustion. Instead, our data suggest B-cell depletion and NK cell dysfunction could be alternative pathways for immune dysregulation in EAC with LOY, implying that tumor cells with LOY can evade immune surveillance via mechanisms other than T-cell exhaustion.

EAC samples from patients with obesity had fewer cytotoxic and regulatory T cells and B cells of various types than those from patients who were normal weight (Figs. 4 I–J). Age may partially account for these differences (Fig. S18). A similar “immune-effector desert” has been described for other obesity-associated cancers, wherein cytotoxic T lymphocyte programs and B-cell antigen-presentation modules collapse under metabolic stress(*28, 29*). Our data are consistent with the “obesity paradox,” where obesity leads to leptin- or PD-1–driven T-cell dysfunction, renewing tumor sensitivity to the PD-1/PD-L1 blockade. Clinically, obese EAC patients could potentially benefit from checkpoint inhibitors paired with agents, such as cytokines, that enhance cytotoxic T lymphocyte recruitment to make tumors more responsive to treatment(*30*). The effects of these demographic and clinical variables when coupled with gene expression changes highlight the complex influences driving EAC tumor evolution, including key transcriptional changes in cell subpopulations and the roles of genetic and environmental factors in shaping the EAC TME.

### LOY and X-chromosome Duplication is Common in EAC and BE Cell Lines

To investigate if LOY is common in BE and EAC cell lines, we performed shallow whole-genome sequencing on one male-derived (OE19) and one female-derived (OE33) EAC cell line, four male-derived BE cell lines (CP-A, -B, -C, and -D) (Fig. S20). Notably, four of the male-derived cell lines exhibited moderate to complete LOY (three BE, one EAC), but the male-derived non-dysplastic BE cell line (CP-A) had almost no LOY. The extent of LOY appeared to correlate with disease progression (Figs. 2K, S20). Full or partial X-chromosome duplication was seen in the CP-D and OE19 cell lines, mirroring observations from patient samples (Fig. 2).

### Cells with LOY Outcompete Those with Intact Y Chromosomes in BE and Colon Cancer Cell Lines

To explore the dynamics of LOY within cell populations, we performed shallow whole-genome sequencing on two human cell lines, HCT116 (colon cancer, all EAC male cell lines have complete LOY) and CP-B (BE, pre-EAC state), across multiple passages (Material and Methods). Both lines showed escalating LOY (Fig. 5), while the X chromosome remained relatively stable (Fig. 5). This progressive reduction in Y chromosome CN (Figs. 5A-H, slope: – 0.008 copies per cell division) suggests that cells with LOY could outcompete cells with an intact Y chromosome, possibly due to inactivation of Y-linked tumor suppressor genes or other regulatory elements. In HCT116 lines, an expanded view of the CN analyses (Figs. 5I–L) confirmed the complete LOY. In addition, we identified 1% dimethyl sulfoxide (DMSO) treatment significantly reduce the speed of LOY by (Figs. 5M-N, *p*= 0.025, paired, two-tailed t-test). A similar but more modest trend was observed in CP-B (Figs. 5O–R, slope: –0.004 copies per cell division), potentially reflecting its origin as a precancerous dysplasia from a BE patient. The CP-A line (from BE with metaplasia) did not lose copies of the Y chromosome over time (Fig. S21), despite the serial passage methods, culturing conditions, and medium being identical to those for CP-B cells (methods). Altogether, we conclude that oxidative damage underlies the observed LOY. These observations support the hypothesis that LOY confers a survival or proliferative advantage and may therefore contribute to cancer evolution in male patients. Consistent with this oxidative-stress model, DMSO, a strong reactive oxygen species scavenger, has been reported to increase lifespan in model organisms(*31*).

**Fig. 5.**
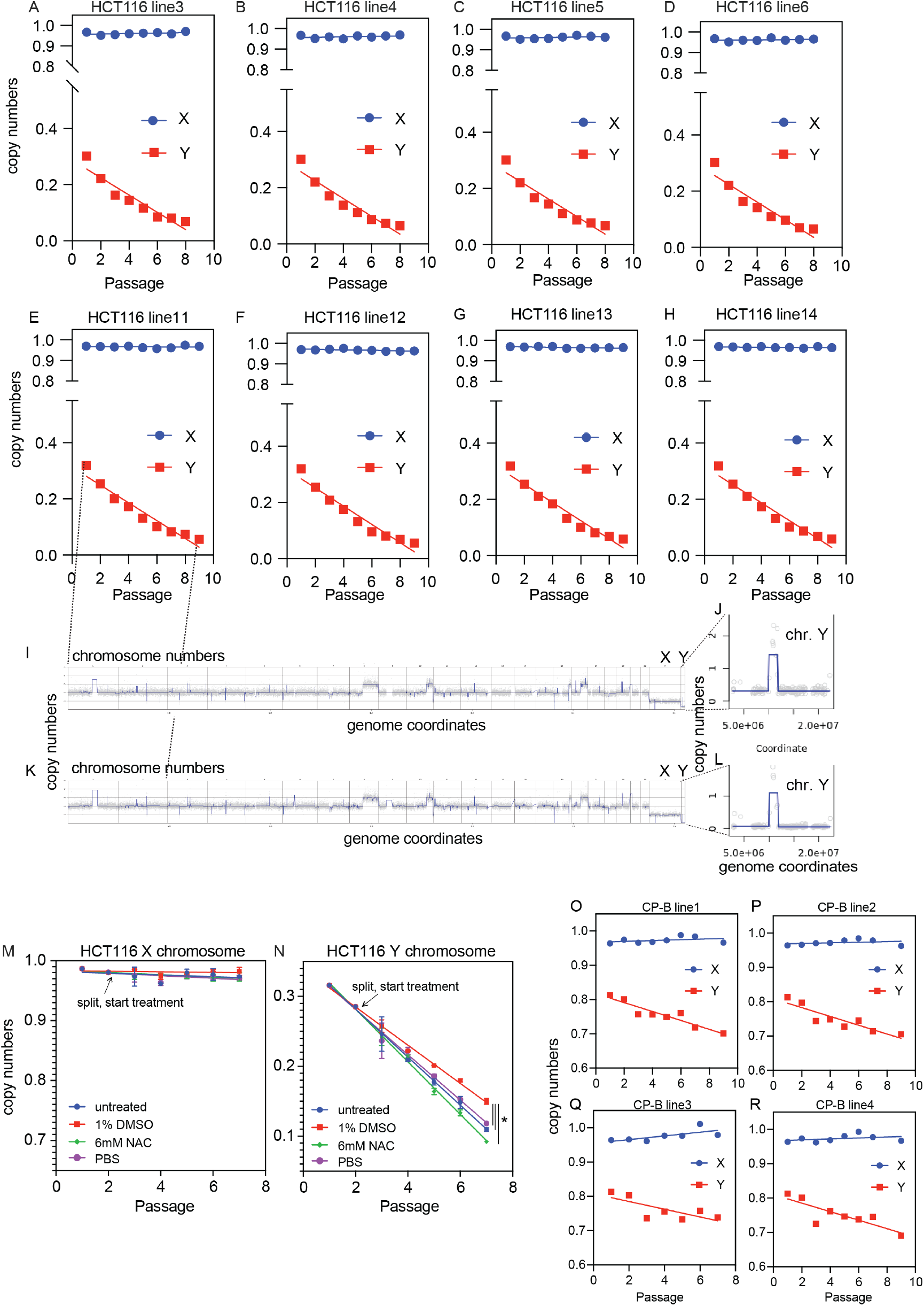
Cell Line Models to study LOY mechanisms and prevention. (A–L) Shallow whole-genome sequencing data for HCT116 cell lines across different passages. Blue circles represent X-chromosome copy numbers, and red squares depict Y-chromosome copy numbers. The x-axis shows the passage number (1–9), and the y-axis indicates copy numbers. Panels A–H show progressive LOY across different HCT116 cell lines, while the X chromosome copy numbers remain stable. Panels I–L show the representative copy number of the whole genome for one of the HCT116 line11; panels J and L are expanded views of the Y chromosome only. (M-N) 1% DMSO significantly reduces speed of LOY. (O–R) Shallow whole-genome sequencing data for CP-B cell lines showing similar LOY and stable X-chromosome copy numbers across passages.

## Discussion

Our single-cell analysis revealed LOY as a frequent and functionally significant genomic alterations in EAC. LOY occurs at a higher rate in EAC than in most other solid tumors(*22*), positioning it as a potential complimentary or substitute molecular marker that does not rely on endoscopic technique and biopsy sampling, like dysplasia. Because LOY is also linked to an elevated neoantigen burden, lesions with LOY could be inherently more immunogenic. Taken together, these observations suggest that LOY status could support early intervention with local ablative technologies and inform targeted immunotherapy-based treatment strategies.

From our comparison of tumor tissues and peripheral blood mononuclear cells, LOY appears to be a somatic event in EAC, given the stability of the Y chromosome in non-tumor blood cells compared with tumor cells. Tumor-specific LOY often co-occurs with complex somatic mutations (Figs. 2B-C, E-G), suggesting that LOY confers a selective advantage in the TME. Nevertheless, the mechanisms through which LOY supports tumorigenesis is poorly understood. Several Y-linked genes, including *UTY, KDM5D*, are implicated in tumor suppression or immune modulation(*10*). *UTY* encodes a histone demethylase that helps regulate gene expression and proliferation(*32*), and *KDM5D* contributes to chromatin remodeling(*32*). Losing these genes could permit unchecked tumor growth or immune evasion, especially if LOY eliminates critical regulatory functions.

In our study, LOY frequently coincided with X-chromosome duplication; these cells had an XX-like genotype without X inactivation. This duplication could potentially upregulate certain X-linked genes, conferring additional selective advantages, such as immune evasion or adaptation to oncogenic stress; further investigation is needed to confirm. Our snRNA-seq revealed associations between LOY status, the immune microenvironment, and patient characteristics, such as obesity. Although obesity typically correlates with an immunosuppressive profile (*33*), patients with obesity can paradoxically respond favorably to immune-checkpoint blockade (*34, 35*). This apparent contradiction warrants examining if LOY amplifies or attenuates the tumor’s immunosuppressive milieu, which could inform the selection of therapies that boost antitumor immune activity, such as bispecific T-cell engagers, oncolytic viral vectors, therapeutic vaccines, and other agents that heighten immunogenicity.

This study has several limitations. First, the small discovery cohort limits statistical power. Second, the low number of paired pre- and post-treatment samples precludes determining if treatment induces or selects for LOY. However, rat models of EAC (e.g. reflux-induced model via esophagogastroduodenal anatomosis) (*36*) could serve as a suitable in vivo system for further interrogating mechanisms and future cancer-prevention strategies.

In summary, LOY is a somatic, potentially actionable genetic alteration in male patients with EAC. LOY could potentially serve as a marker for early detection, risk stratification, and immunotherapeutic response, particularly in patients with obesity. We acknowledge certain limitations, including a modest cohort size and the possibility that LOY in adjacent squamous mucosa could arise from prior treatment rather than being a BE-independent mechanism for developing EAC. Prospective and pan-cancer investigations coupled with further mechanistic studies are needed to validate these findings and clarify how LOY, X-chromosome duplication, and co-occurring mutations influence EAC progression. By integrating broader molecular analyses with key clinical variables, like obesity, future work can further unveil the complexities of LOY in EAC and guide personalized disease management.

## Acknowledgments

We thank the Creighton University Innovative Genomics and Histology Cores for their assistance; Yangzi Liu for experimental support; the research coordinators in the Creighton Clinical Research Office for their efforts; Dr. Peter S. Rabinovitch for generously providing the CP-A, -B, -C, and -D cell lines via Dr. Kresty; and Dr. Katherine H. Sippel for her expert editorial assistance.

## Funding

State of Nebraska LB595 grant (JX, YF)

State of Nebraska LB692 grant (JX)

Kicks for a Cure foundation (JX, YF, KCN, WBA)

National Institutes of Health grant R00ES033259 (JX)

Texas A&M HSC startup fund (JX, YF)

Creighton University Surgery Department Chair Fund (WBA)

## Author contributions

Conceptualization: JX, YF, KCN

Methodology: YF, JX, KCN, SY, NZ, BF, SP, NKJ, SA, NV

Pathology examination: ND

Investigation: all authors

Visualization: JX, YF

Funding acquisition: JX, YF, WBA, KCN

Project administration: JX, YF, KCN, WBA

Supervision: JX, YF, KCN

Writing – JX, YF, KCN, EAG, WBA

Materials: KCN, SKP, LK, JX, YF

Writing – review & editing: all authors

## Data and Materials Availability

All data required to evaluate the conclusions of this study are provided in the main text and supplementary files. Cell lines are available from ATCC or Sigma. Due to Dr. Rabinovitch’s retirement, CP-A, -B, -C, -D lines are more easily assessable from ATCC.

## Supplementary Materials

Materials and Methods

Figs. S1 to S21

Tables S1 to S3

References (*37–45*)

