## Supplementary material for "Single-cell Genomic Copy Number Evolution Reveals Frequent Loss of the Y chromosome in Esophageal Adenocarcinoma"

**The PDF file includes:**

Materials and Methods
Figs. S1 to S21
Tables S1 to S3

### Materials and Methods

#### Study subjects, sample collection and storage

This study was conducted after receiving approval from the Institutional Review Board (IRB) of Creighton University (IRB No. 1194896). It also received a Human Research, non-engaged determination from Texas A&M (IRB No. STUDY2024-0977). Patients were recruited from the surgery clinics at Creighton University Medical Center and CHI Health Immanuel Medical Center, and informed consent was obtained from all participants. All participants include clinical diagnosis of Barrett's esophagus or EAC confirmed by endoscopic inspection of the esophagus. Normal tissue samples and PBMC were also collected for most participants. Exclusion criteria included those ages 18 years or younger, individuals unwilling to participate in the study. Detailed information on participants and data availability from this study, see Table S1.

Esophageal tissue biopsies—from both tumor and adjacent non-tumor regions—as well as peripheral blood were obtained during endoscopy or surgery. All specimens were immediately transported on ice to the Creighton University laboratory and stored at –80 °C until processing. Single-nucleus or single-cell suspensions were generated using the gentleMACS Octo dissociator with heaters (see "Single-cell/nucleus isolation" below), followed by 10× Genomics single-nucleus or single-cell RNA-seq and/or single-cell/nucleus copy-number profiling. In total, 20 specimens yielded single-cell DNA-sequencing data, and 32 specimens underwent single-nucleus RNA-seq; these included normal esophageal lining, Barrett's esophagus, and esophageal adenocarcinoma (EAC) samples collected from 17 consented patients. Additionally, we performed whole-genome sequencing on 17 paired PBMC samples.

For each EAC case, a ~2 mm biopsy was collected from the lesion, and a matched control biopsy of similar size was taken from adjacent normal tissue. Participants provided informed consent and completed demographic, medical (including weight and BMI), surgical, and social history questionnaires before their first visit. On the day of surgery or endoscopy (second visit), 20 mL of blood was drawn, and standard-of-care biopsies were obtained from both diseased and normal esophageal tissue. A third visit applied to patients receiving neoadjuvant chemotherapy: these individuals donated an additional 5 mL of blood and underwent follow-up biopsy at the time of surgery.

#### Single nucleus isolation

Single nuclei were isolated from tissue by performing mechanical and enzymatic dissociation. Up to 50 mg of tissue was minced into small fragments and then incubated in 4 ml of Nuclei Extraction Buffer (Miltenyi Biotec) combined with 20 µl of RNase inhibitor buffer (Invitrogen, ThermoFisher). This mixture was transferred to a gentleMACS C Tube with ice-cold lysis buffer and subjected to dissociation in a gentleMACS Octo Dissociator at 4°C for 5 minutes. The dissociation step was repeated to ensure thorough breakdown of tissue fragments. The resulting sample was filtered through a 70 µm MACS strainer, diluted with 1 ml lysis buffer, and then centrifuged at 300 × g for 5 minutes. The supernatant was removed, and the cell pellet was resuspended in 1 ml lysis buffer; a further 1 ml was used to rinse the tube. The sample was subsequently passed through a 30 µm MACS strainer and again centrifuged at 300 × g for 5 minutes. After centrifugation, 200 µl of PBS was added to the pellet, and a 20 µl aliquot was taken for counting. These suspensions were used for downstream 10x single-cell RNA-seq pipeline right away onto a 10X Gem-X 3' or 5' chip and processed on the 10x Chromium X/iX. For single-nucleus DNA copy number pipelines, 1 ml of PBS was added to both stained (with Vybrant™ DyeCycle™ Green Stain, V35004) and unstained samples to adjust the total volume. Single-nucleus sorting was performed using either a BioRad S3e cell sorter (into 8-strip tubes) or a BD Aria II cell sorter (into 96-well plates). Each well (or tube) was pre-filled with 2 µl of lysis buffer (30mM Tris-HCL, pH8.0, 10mM NaCl, 5mM EDTA, 0.2% Triton, proteinase K 2mg/ml) After sorting, plates and tubes were spun at 2000 rpm for 2 minutes at 4°C, then incubated at 65°C for 3 hours to facilitate cell lysis, followed by another spin at 2000 rpm for 2 minutes at 4°C. Samples were then stored at −80°C for further copy number analyses.

**Single-cell whole genome amplification and sequencing**

Single cells (or single nuclei) in 96-well plates were lysed (described above), then briefly centrifuged to the bottom. For tagmentation, each well was brought to 10 µL with a mixture containing 1× TD buffer, 0.01 µL TTE Mix V50 (Vazyme TD501), 0.625× protease inhibitor cocktail (Promega G6521), and 1 mM $MgCl_2$. The plate was incubated at 55°C for 2 hours. After tagmentation, 12 µL of PCR master mix was added to each well, consisting of 11 µL Q5 High-Fidelity 2× Master Mix (M0492S, New England Biolabs) and 0.5 µL each of i5 and i7 index primers [see Table S3,(*37*)]. The PCR program was 72°C for 5mins, 98°C for 30 seconds, followed by 24 cycles of 98°C for 15 seconds, 60°C for 30 seconds, and 72°C for 90 seconds, with a final extension at 72°C for 5 minutes. PCR products from multiple plates were pooled and purified using DNA Clean & Concentrator kits (Zymo). Quality control was performed using an Agilent TapeStation 4200. Final libraries were sequenced on an Illumina NextSeq 2000 System using paired-end 2×61 bp reads with an 8 bp + 8 bp dual index.

**Hematoxylin and eosin (H&E) staining**

Tissue-staining protocols were described previously(*38*). In brief, samples for histological analysis (Figs. S12-13) were fixed in formalin for 14–18 h, dehydrated in 70 % ethanol overnight, and then processed for automated hematoxylin and eosin (H&E) staining. Stained slides were scanned on an Olympus VS120 Slide Scanner, first at low resolution (4×) for a quick overview, and then at high resolution (20×) to capture detailed tissue morphology. All H&E slides were reviewed by a board-certified pathologist.

**Fluorescence in situ hybridization (FISH)**

Fluorescence in Situ Hybridization (FISH) was used to detect the loss of the Y chromosome and to quantify chromosome 1, X, and Y copy numbers. For the quantification of chromosome 1, X, and Y copy numbers in formalin-fixed, paraffin-embedded (FFPE) tissues, sections were deparaffinized and fixed with 4% paraformaldehyde/phosphate-buffered saline (PFA/PBS) for 30 minutes at room temperature. The sections were then incubated with 0.004% pepsin in 0.1 N HCl at 37°C for 5 minutes. A mixture of 10 µl Human XY Chromosome FISH Probe (Cy3 and Cy5, FHXY-01, Creative Bioarray) and 1 µl Human CEN 1p FISH Probe (FITC, FCEN-01p, Creative Bioarray) was applied to each section, followed by denaturation at 90°C for 10 minutes and overnight hybridization at 37°C. After post-hybridization washes, the slides were counterstained with DAPI, mounted with an anti-fade medium, and imaged using the CW4000 FISH application program (Leica Microsystems Imaging Solution Ltd.) on a Leica DMRA2 microscope equipped with a cooled CCD camera.

**Principal component analysis for cell of origin**

Single nucleotide polymorphisms (SNPs) were called directly from shallow single-cell whole-genome sequencing (scWGS) data using Samtools/Bcftools. Although shallow scWGS yields sparse coverage (primarily for copy number analysis), SNP calls from all single cells in each sample group (e.g., P2EAC x=1 and P2EAC x=2) were aggregated to create a combined genotype profile. These aggregated genotypes (one per group) were compiled into a matrix with samples as rows and SNP loci as columns. Principal component analysis (PCA) was then performed on this matrix to reduce dimensionality and assess genetic similarity among samples. The resulting principal components were examined to verify that x duplicated cell groups clustered with their corresponding non-duplicated samples, thereby confirming that both cell sets originated from the same individual.

**10x Genomics single-nucleus RNA-seq**

Single-nucleus RNA sequencing (snRNA-seq) libraries were prepared using the Chromium Next GEM Single Cell 3′ v3.1 or Chromium GEM-X Single Cell 3′ v4 Gene Expression kits from 10x Genomics, following the manufacturer's protocols. For the v3.1 protocol, the Chromium Next GEM Single Cell 3′ Kit v3.1 (PN-1000121) and Chromium Next GEM Chip G Single Cell Kit (PN-1000120) were used. For the v4 protocol, the Chromium GEM-X Single Cell 3′ Kit v4 (PN-1000779) and Chromium GEM-X Single Cell 3′ Chip Kit v4 (PN-1000747) were used. In both protocols, sample indexing was performed using the Dual Index Kit TT Set A (PN-1000215). Briefly, single-cell suspensions were loaded onto the Chromium Controller to generate Gel Beads-in-Emulsion (GEMs), enabling cell barcoding and reverse transcription for cDNA synthesis. Following post-GEM-RT cleanup and cDNA amplification, library construction involved fragmentation, end repair, A-tailing, adapter ligation, and sample index PCR. Final libraries were quality-checked on an Agilent TapeStation 4200 and quantified with a Qubit Fluorometer (Thermo Fisher Scientific), pooled to 650 pM, and sequenced on an Illumina NextSeq 2000 with paired-end reads (Read 1: 28 bp; Read 2: 90 bp; dual indices: 10 bp each) per 10x Genomics recommendations.

**Single-nucleus RNA-seq data analysis**

We processed these fastq files using Cell Ranger (v6.1.2; 10x Genomics) with the human GRCh38 reference, which performed demultiplexing, alignment, and quantification to generate filtered gene-barcode matrices. Downstream analyses were carried out in R (v4.3) with Bioconductor (v3.2) and in Python using Scanpy(*39*). Cells with fewer than 300 detected genes, over 30% mitochondrial transcripts, or predicted as doublets by Scrublet (v0.2)(*40*) were removed. The data were then normalized to 10,000 counts per cell, log-transformed, and filtered for highly variable genes (e.g., min_mean=0.0125, max_mean=3, min_disp=0.5). To correct for sample batch effects, we applied BBKNN (*41*) (using 50 principal components) before computing a k-nearest neighbor graph (k=15) for dimensionality reduction by PCA and UMAP. We clustered the cells using the Leiden algorithm (resolution=3.0), and subsequently annotated clusters with CellTypist(*42*) (Immune_All_Low model) in majority-voting mode, complemented by cello-based annotation and manual evaluation of marker genes.

**Detecting copy number changes in single nucleus and bulk samples**

Single-cell nucleus analytical approached were adapted from (*18, 37*). Raw FASTQ files were demultiplexed and subjected to Fastp for adapter trimming and removal of low-quality reads. Single nuclei with files sizing less than 1Mb were removed. Trimmed reads were aligned to the GRCh38 human reference genome using Bowtie2(*43*). only single nucleus with more than 100k reads and with an overall alignment rate of 50% or higher were included in subsequent heatmap analyses. SAM files were binned using 50kb and 500kb bins to facilitate coverage-based assessments, then converted to sorted and indexed BAM files using SAMtools(*44*). Copy number alterations were identified using a single circular binary segmentation (CBS) approach(*45*), which segments the genome into regions of distinct copy number. Resulting profiles were examined to detect amplifications and deletions in single cells, enabling the identification of copy number alterations from individual nucleus. Bulk NEB samples from blood or cell lines samples were using the same pipeline for copy number analysis.

**Genomic DNA extraction from blood**

Genomic DNA was extracted from blood samples using the QIAGEN DNeasy Blood & Tissue Kit, following the manufacturer's instructions. Briefly, 50–100 µl of anticoagulant-treated blood was mixed with 20 µl of Proteinase K and adjusted to a total volume of 220 µl with phosphate-buffered saline (PBS). After adding 200 µl of Buffer AL, the mixture was incubated at 56°C for 10 minutes to ensure complete lysis. Subsequent steps included the addition of ethanol, transfer to a DNeasy Mini spin column, washing with Buffers AW1 and AW2, and elution of DNA with Buffer AE. The concentration and purity of the extracted DNA were assessed using Qubit Fluorometric Quantification.

**Library preparation using the NEBNext Ultra II FS DNA library prep kit**

Genomic DNA (gDNA) extracted from blood samples and cell lines was used to construct sequencing libraries with the NEBNext Ultra II FS DNA Library Prep Kit for Illumina (NEB #E7805). For each library, 100 ng of gDNA was processed following the manufacturer's protocol. The fragmentation, end repair, and dA-tailing steps were performed in a single reaction by adding 7 µl of NEBNext Ultra II FS Reaction Buffer and 2 µl of NEBNext Ultra II FS Enzyme Mix to the 100 ng DNA sample, resulting in a total volume of 35 µl. The reaction was incubated at 37°C for 15 minutes, followed by heat inactivation at 65°C for 30 minutes. Subsequent steps included adaptor ligation, USER enzyme treatment, size selection using SPRIselect beads, and PCR amplification with NEBNext Ultra II Q5 Master Mix and index primers. Library quality was assessed using a Bioanalyzer, Qubit, and TapeStation prior to sequencing.

**Library preparation using the FS Pro DNA Lib Prep kit**

Genomic DNA (100 ng) isolated from human blood samples and cultured cell lines was used for library construction with the FS Pro DNA Library Prep Kit V2 (ABclonal, Cat. No. RK20275). DNA fragmentation, end repair, and dA-tailing were carried out in a single reaction by combining the template with 5 µL FS Pro Buffer I and 13 µL FS Pro Enzymes II in a 50 µL reaction volume. Samples were incubated at 32 °C for 10 min, followed by 72 °C for 30 min. Adapter ligation was performed by adding 20 µL FS Pro Ligation Buffer II, 5 µL Ligase Enzymes, and 5 µL universal adapter, and incubating at 22 °C for 15 min. Ligation products were purified using SPRI beads (Beckman Coulter, Cat. No. B23318) at a 0.8× bead-to-sample ratio and eluted in 20 µL nuclease-free water. Libraries were amplified with the 2× PCR Mix and 5 µL of Unique Dual Index primers (ABclonal, RK21624 or RK21625) using six PCR cycles. Final products were purified with 1× SPRI beads and eluted in 30 µL nuclease-free water. Library concentration and fragment size distribution were assessed using a Qubit fluorometer and TapeStation system prior to sequencing.

**HCT116 cell culture conditions and anti-LOY drug screening**

HCT116 colon cancer cells (ATCC CCL-247) were obtained from ATCC and authenticated by shallow whole-genome sequencing. Cells were cultured in Gibco Dulbecco's Modified Eagle Medium (DMEM) supplemented with 10% fetal bovine serum (FBS) and 1% penicillin-streptomycin-glutamine (PSQ). Upon thawing, $1 \times 10^6$ cells were immediately harvested for DNA extraction, while the remaining cells were seeded onto 6-well plates and grown to ~90% confluence over several days. Cells were subsequently passaged every four days at a 1:16 dilution into separate 6-well plates. Genomic DNA (gDNA) was extracted every other passage for each line. DNA concentrations were determined using Qubit™ Fluorometric Quantification, and libraries were prepared with the NEBNext Ultra II FS DNA Library Prep Kit for Fig. 5A–H.

For drug treatment, parental cells were first recovered and harvested for gDNA extraction. Once they reached confluence, additional gDNA was extracted prior to seeding at 1 × $10^5$ cells per well in 6-well plates. Cells were immediately treated in duplicate under one of the following conditions: (i) 1% DMSO (TargetMol, Cat. No. T1662), (ii) 6 mM N-acetyl-L-cysteine (NAC; Sigma-Aldrich, Cat. No. A9165-25G), (iii) 20 µL PBS (Gibco, Cat. No. 14190-250), or (iv) untreated as a negative control. Cells were maintained under standard conditions and passaged every 3–4 days at a 1:16 dilution. At each passage, a portion of the cells was harvested for gDNA extraction using the ABclonal DNAprep Kit (Cat. No. RK20275). DNA concentrations were measured with the Qubit™ Broad Range dsDNA Assay Kit (Thermo Fisher Scientific), following the manufacturer's instructions.

**CP-A, CP-B, CP-C, CP-D cell culture conditions**

CP-A, CP-B, CP-C, and CP-D Barrett's esophagus cell lines (ATCC CRL-4027, CRL-4028, CRL-4029, CRL-4030) were cultured in MCDB 153 medium (Sigma) supplemented with 0.4 µg/ml hydrocortisone, 20 ng/ml recombinant human epidermal growth factor, 8.4 µg/L cholera toxin, 20 mg/L adenine, 140 µg/ml bovine pituitary extract, 1× ITS Supplement (Sigma I1884), 4 mM glutamine, and 5% FBS. They were authenticated via shallow whole-genome sequencing. Thawed cells were initially grown in a T75 flask for seven days until reaching approximately 95% confluence, then split into four T75 flasks at a 1:8 dilution. gDNA extraction occurred every or every other passage. DNA concentrations were determined using Qubit™ Fluorometric Quantification.

**OE19 and OE33 Cell Lines**

Both cell lines are typically maintained in RPMI 1640 medium supplemented with 2 mM glutamine and 10% fetal bovine serum (FBS), cultured at 37°C in a humidified atmosphere containing 5% $CO_2$. Sub-confluent cultures (approximately 70–80%) are usually split at a ratio of 1:8 by seeding cells at 1×$10^4$ cells/$cm^2$ using 0.25% trypsin or trypsin/EDTA. They were also authenticated via shallow whole-genome sequencing. After resuscitation from frozen stocks, these cells often grow in island-like clusters and can take up to seven days to reach about 70% confluence. During this period, partial medium changes (replacing half of the old medium with fresh medium) are typically performed every two to three days to maintain optimal growth conditions.

**Quantification and statistical analysis**

Analytical details can be found in the main text, figure legends and/or material and methods.

**Data and materials availability**

Sequencing data from this study are presented in the paper and/or will be available after publication.

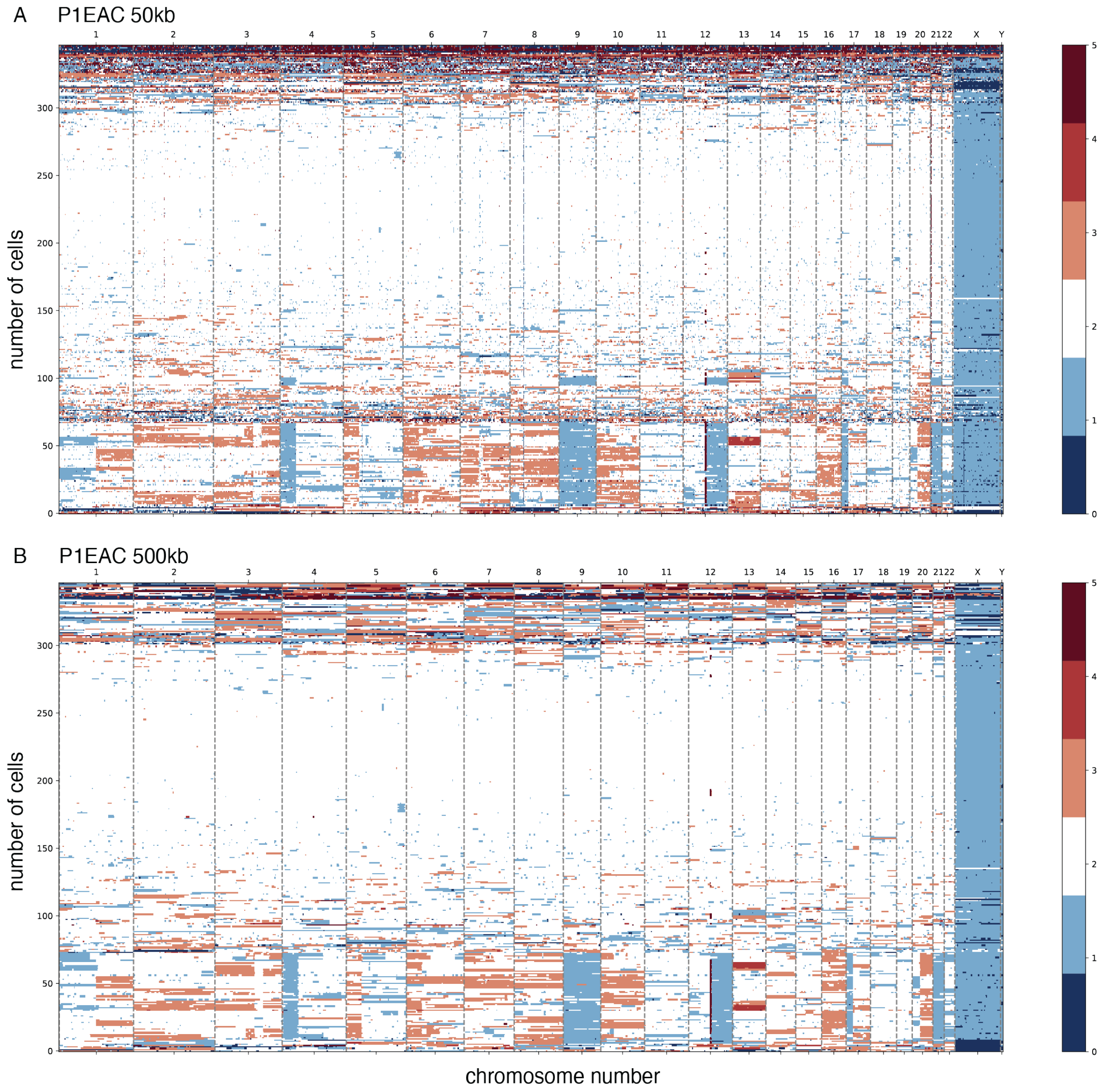


**Fig. S1. Single nucleus Copy Number Heatmap for P1.** EAC: esophageal adenocarcinoma, X axis: genome coordinates, upper: chromosome number. Y axis: cell number. Color scale: copy numbers. White=2, Orange=3, red=4, dark red=5, lighter blue=1, dark blue=0. **A: 50kb bin; B: 500kb bin.**

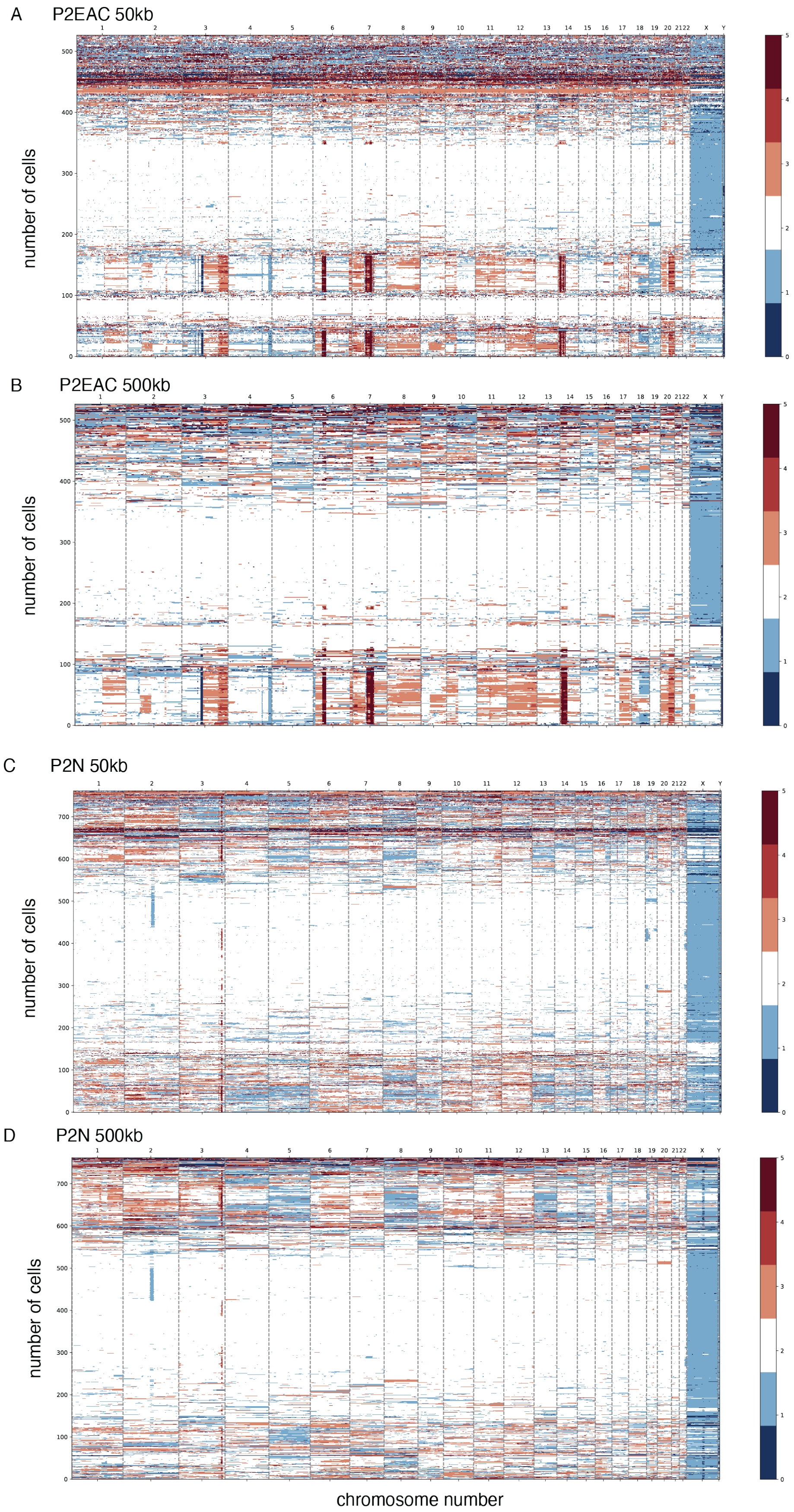


**Fig. S2. Single nucleus Copy Number Heatmap for P2.** EAC: esophageal adenocarcinoma, N: normal esophagus mucosa, X axis: genome coordinates, upper: chromosome number. Y axis: cell number. Color scale: copy numbers. White=2, Orange=3, red=4, dark red=5, lighter blue=1, dark blue=0. **A, C: 50kb bin; B, D: 500kb bin.**

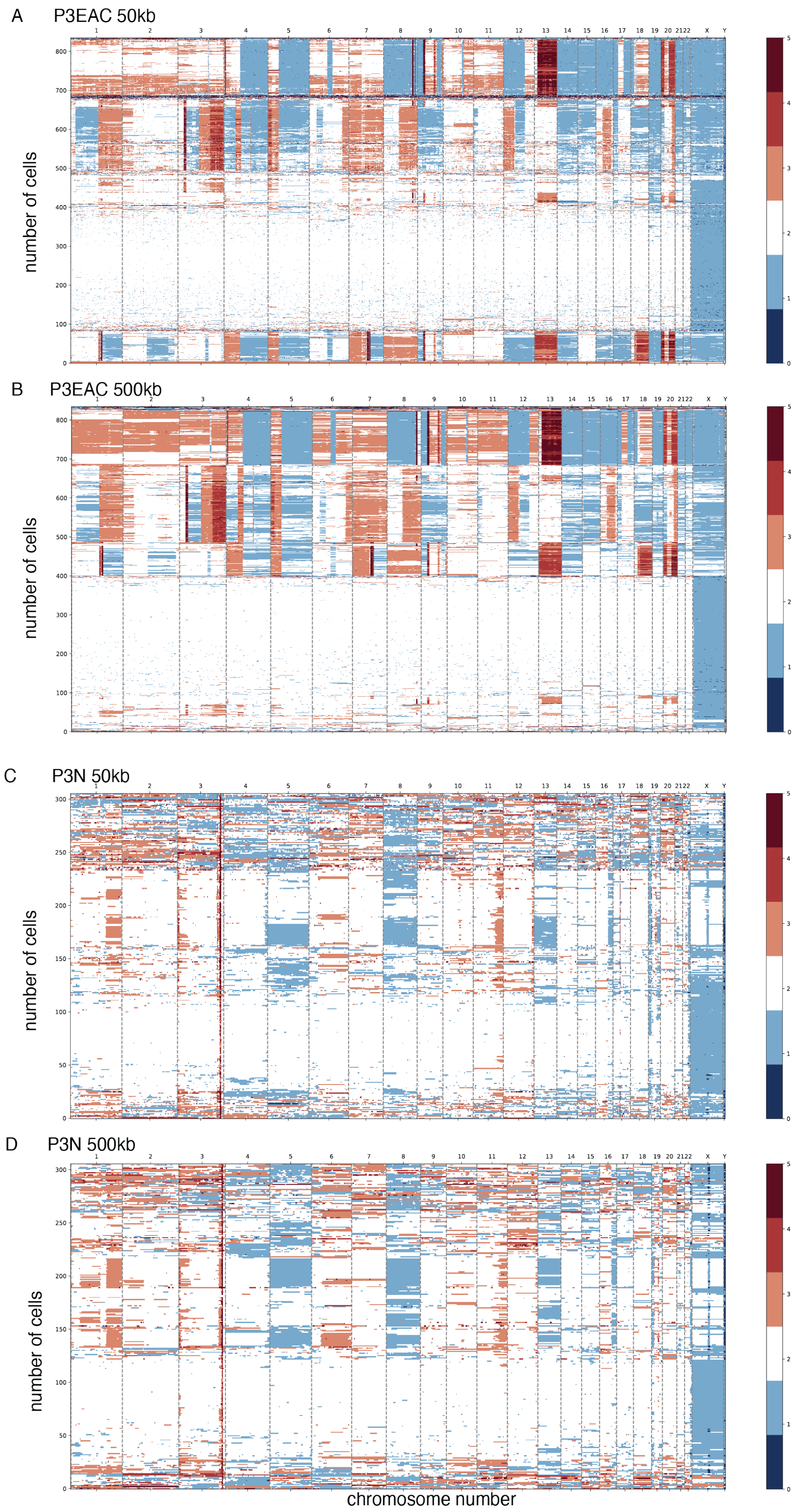


**Fig. S3. Single nucleus Copy Number Heatmap for P3.** EAC: esophageal adenocarcinoma, N: normal esophagus mucosa, X axis: genome coordinates, upper: chromosome number. Y axis: cell number. Color scale: copy numbers. White=2, Orange=3, red=4, dark red=5, lighter blue=1, dark blue=0. **A, C: 50kb bin; B, D: 500kb bin.**

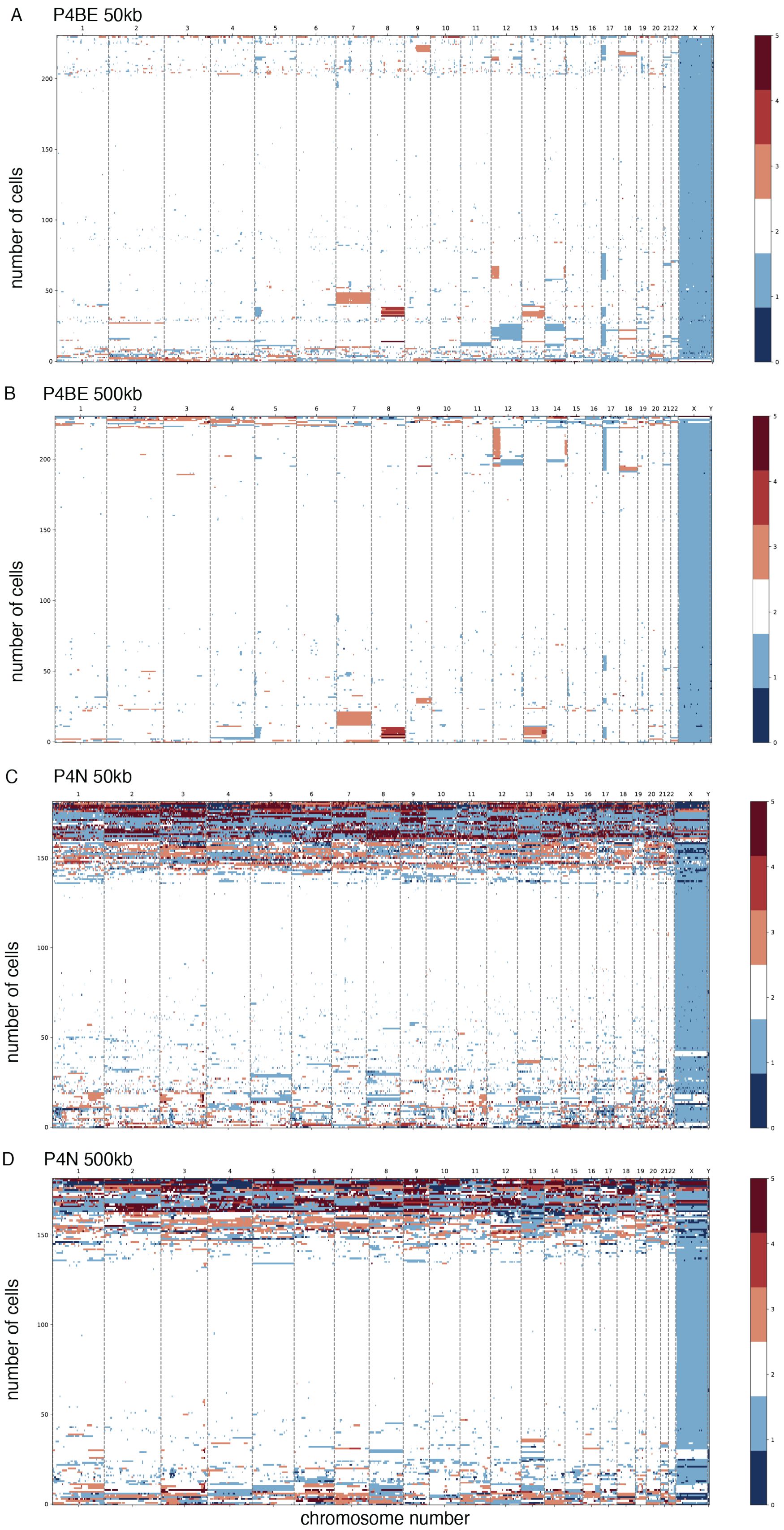


**Fig. S4. Single nucleus Copy Number Heatmap for P4.** BE: Barrett's esophagus. N: normal esophagus mucosa, X axis: genome coordinates, upper: chromosome number. Y axis: cell number. Color scale: copy numbers. White=2, Orange=3, red=4, dark red=5, lighter blue=1, dark blue=0. **A, C: 50kb bin; B, D: 500kb bin.**

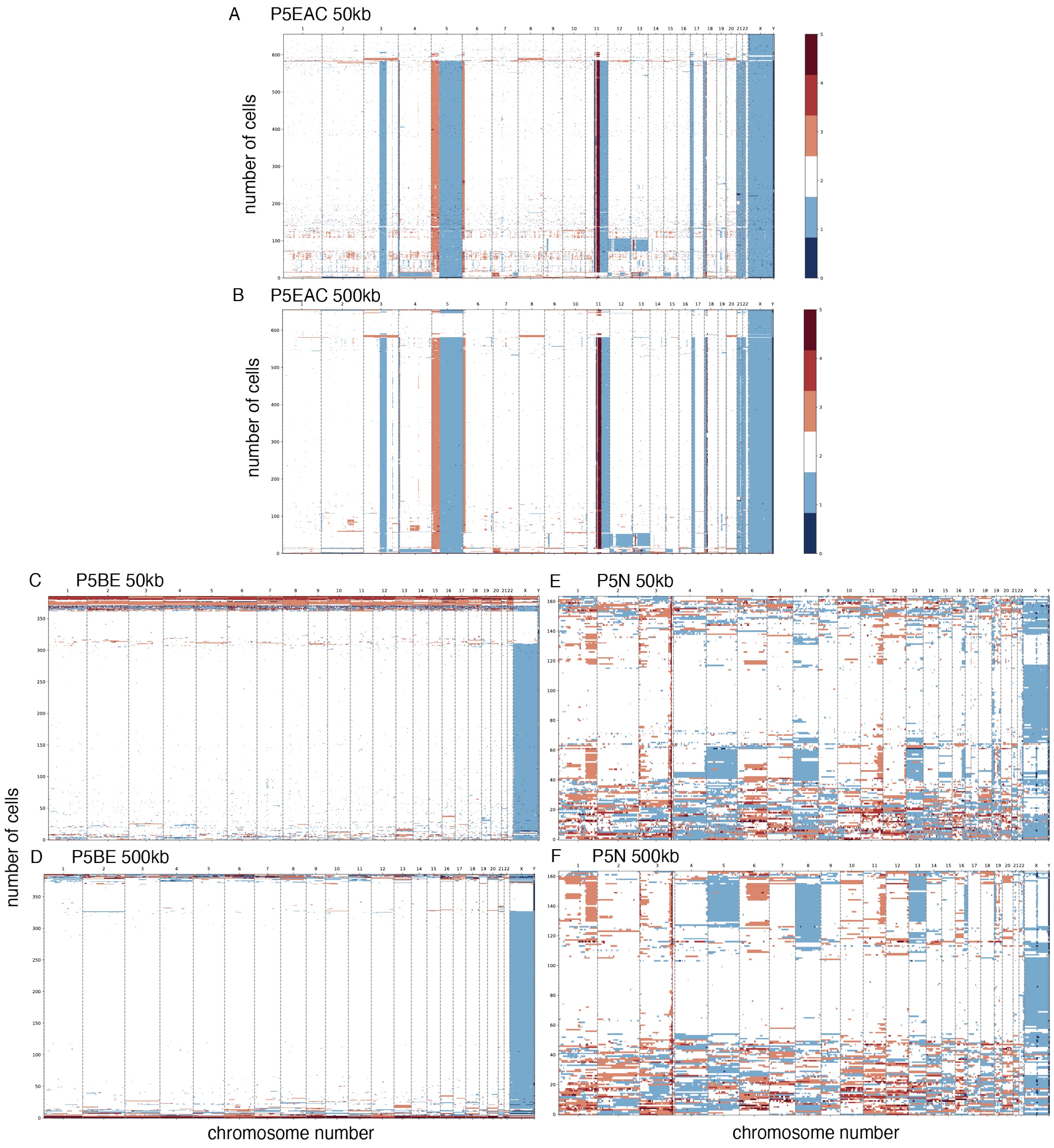


**Fig. S5. Single nucleus Copy Number Heatmap for P5.** EAC: esophageal adenocarcinoma, BE: Barrett's esophagus. N: normal esophagus mucosa, X axis: genome coordinates, upper: chromosome number. Y axis: cell number. Color scale: copy numbers. White=2, Orange=3, red=4, dark red=5, lighter blue=1, dark blue=0. **A, C, E: 50kb bin; B, D, F: 500kb bin.**

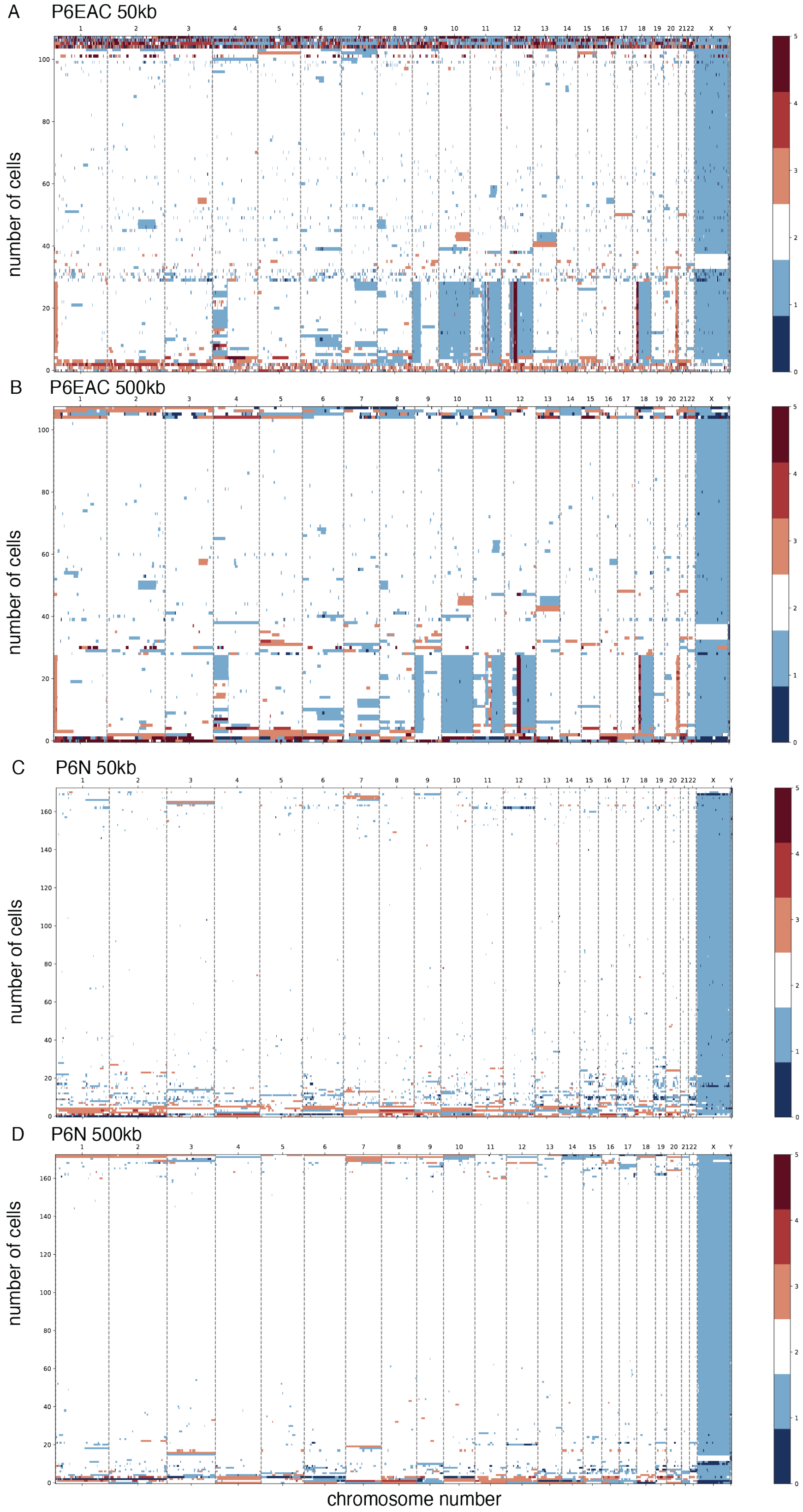


**Fig. S6. Single nucleus Copy Number Heatmap for P6.** EAC: esophageal adenocarcinoma, N: normal esophagus mucosa, X axis: genome coordinates, upper: chromosome number. Y axis: cell number. Color scale: copy numbers. White=2, Orange=3, red=4, dark red=5, lighter blue=1, dark blue=0. **A, C: 50kb bin; B, D: 500kb bin.**

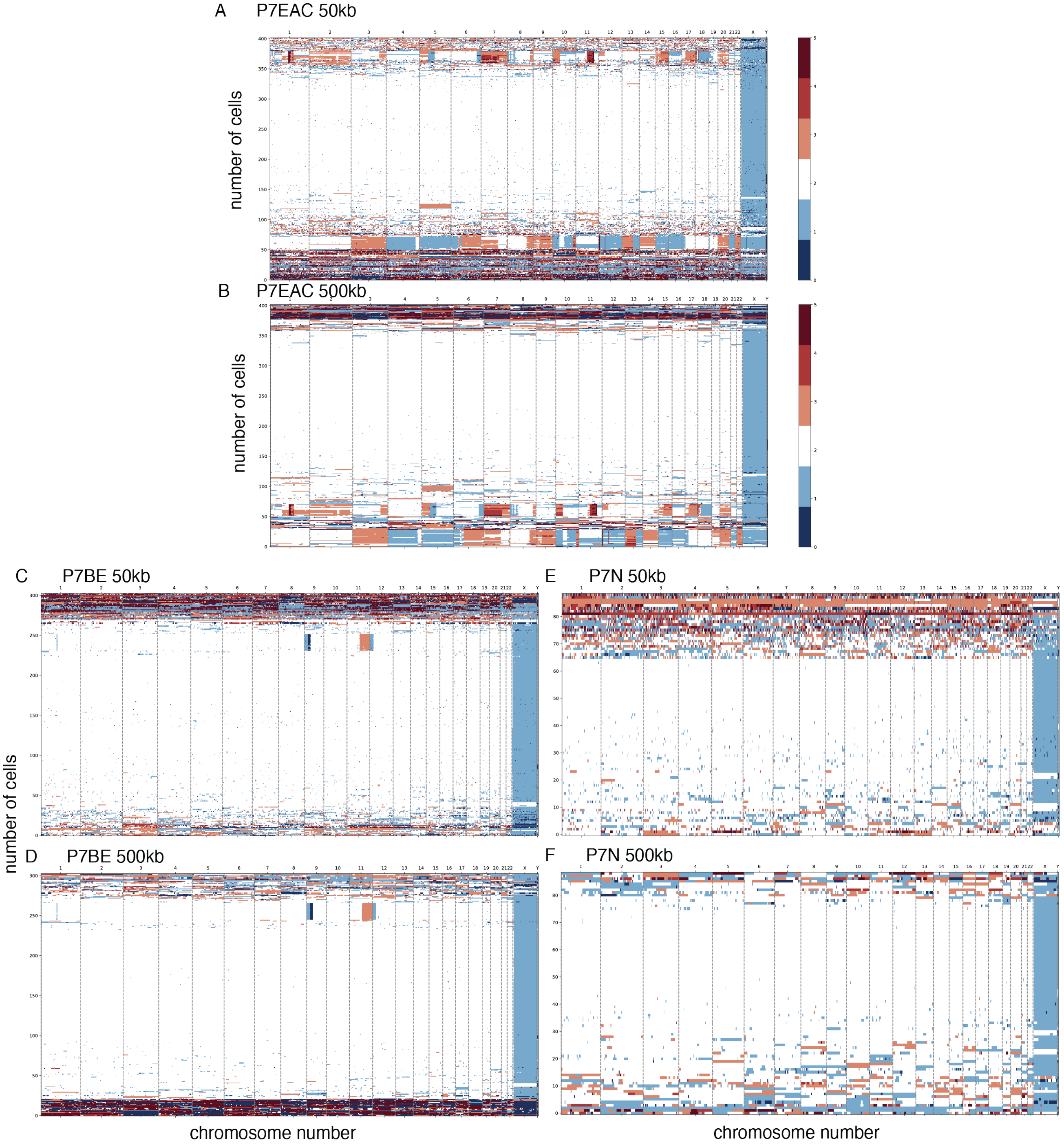


**Fig. S7. Single nucleus Copy Number Heatmap for P7.** EAC: esophageal adenocarcinoma, BE: Barrett's esophagus. N: normal esophagus mucosa, X axis: genome coordinates, upper: chromosome number. Y axis: cell number. Color scale: copy numbers. White=2, Orange=3, red=4, dark red=5, lighter blue=1, dark blue=0. **A, C, E: 50kb bin; B, D, F: 500kb bin.**

5

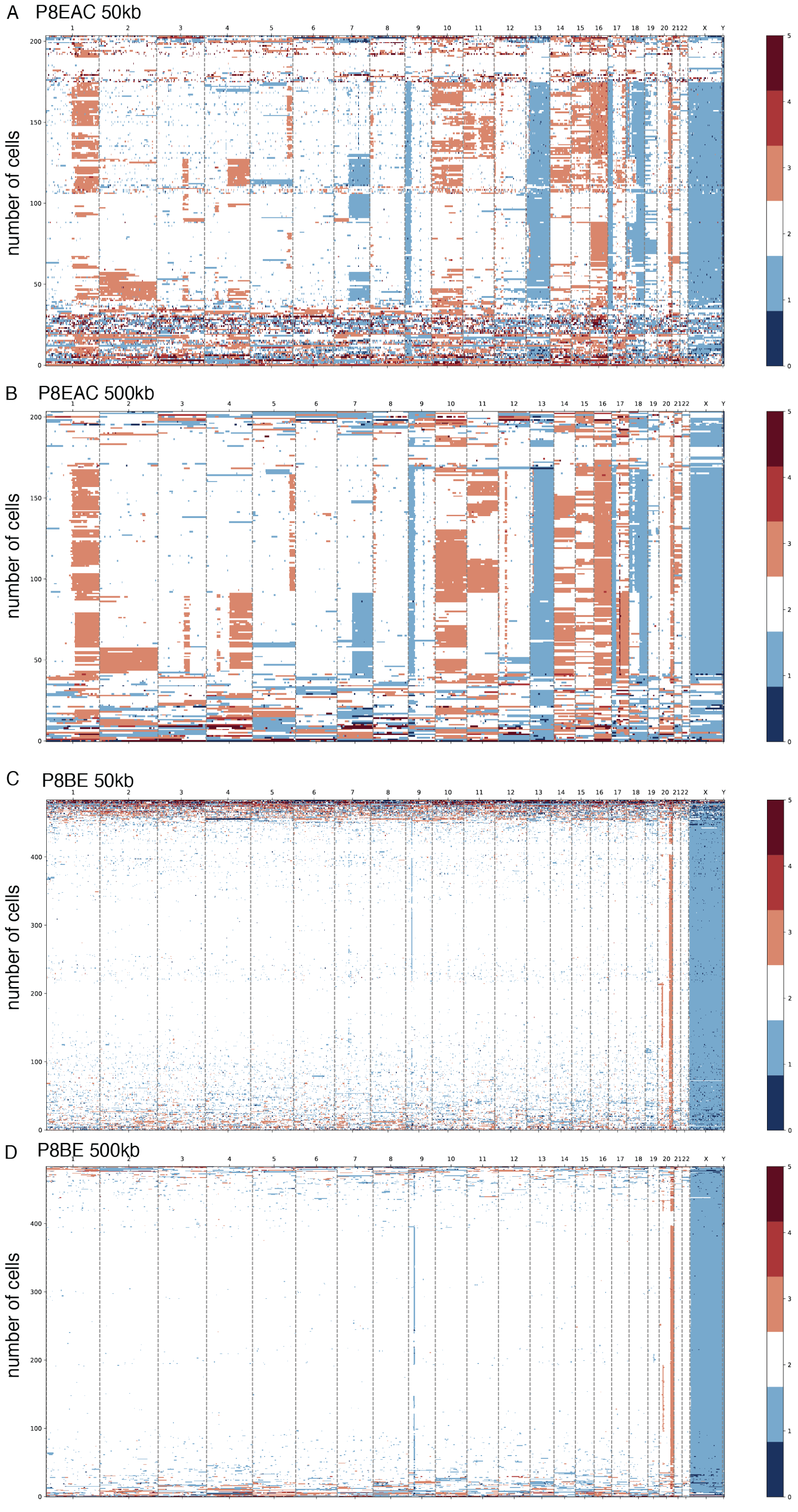


**Fig. S8. Single nucleus Copy Number Heatmap for P8.** EAC: esophageal adenocarcinoma, BE: Barrett's esophagus, X axis: genome coordinates, upper: chromosome number. Y axis: cell number. Color scale: copy numbers. White=2, Orange=3, red=4, dark red=5, lighter blue=1, dark blue=0. **A, C: 50kb bin; B, D: 500kb bin.**

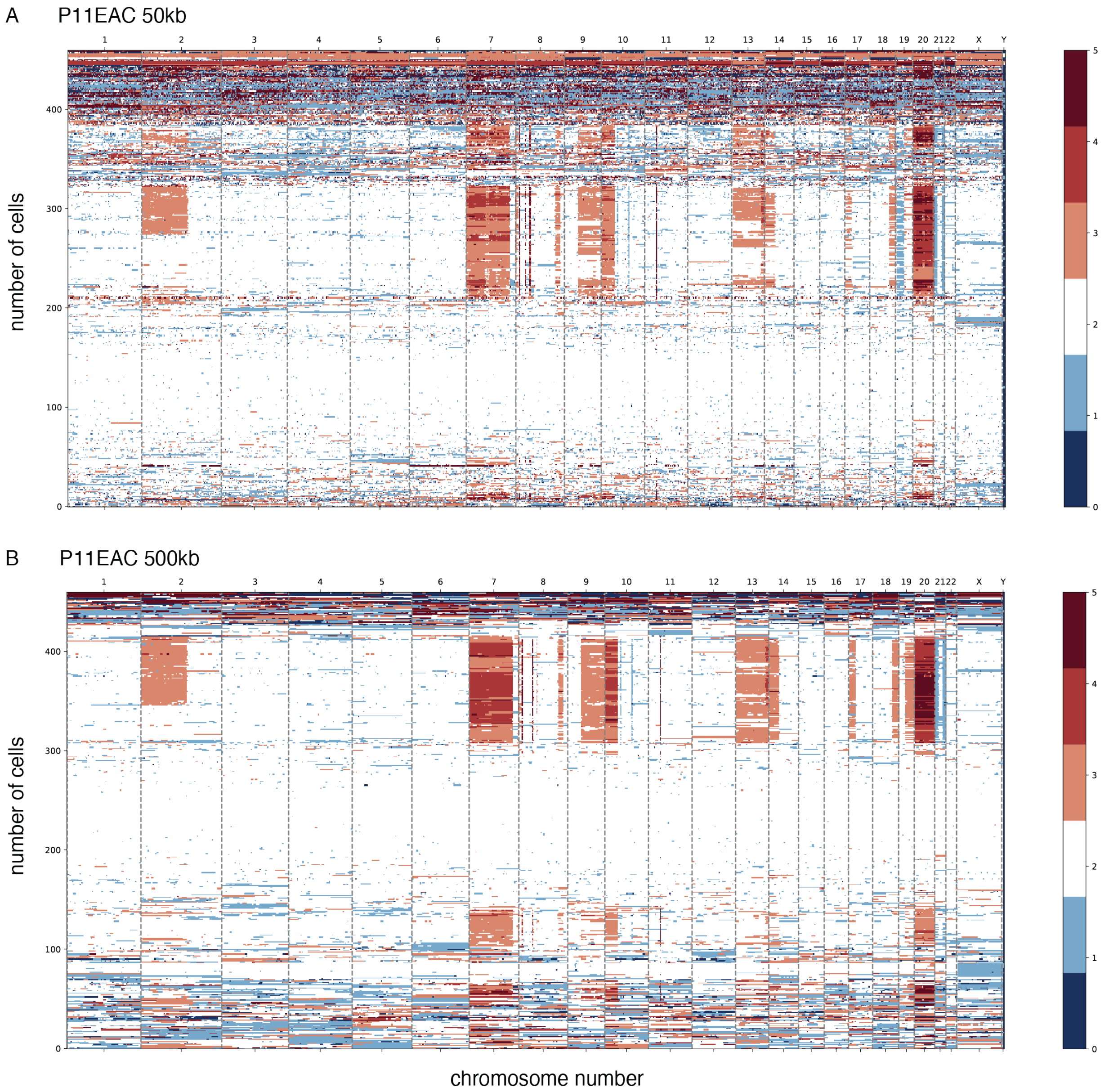


**Fig. S9. Single nucleus Copy Number Heatmap for P11.** EAC: esophageal adenocarcinoma, X axis: genome coordinates, upper: chromosome number. Y axis: cell number. Color scale: copy numbers. White=2, Orange=3, red=4, dark red=5, lighter blue=1, dark blue=0. **A: 50kb bin; B: 500kb bin.**

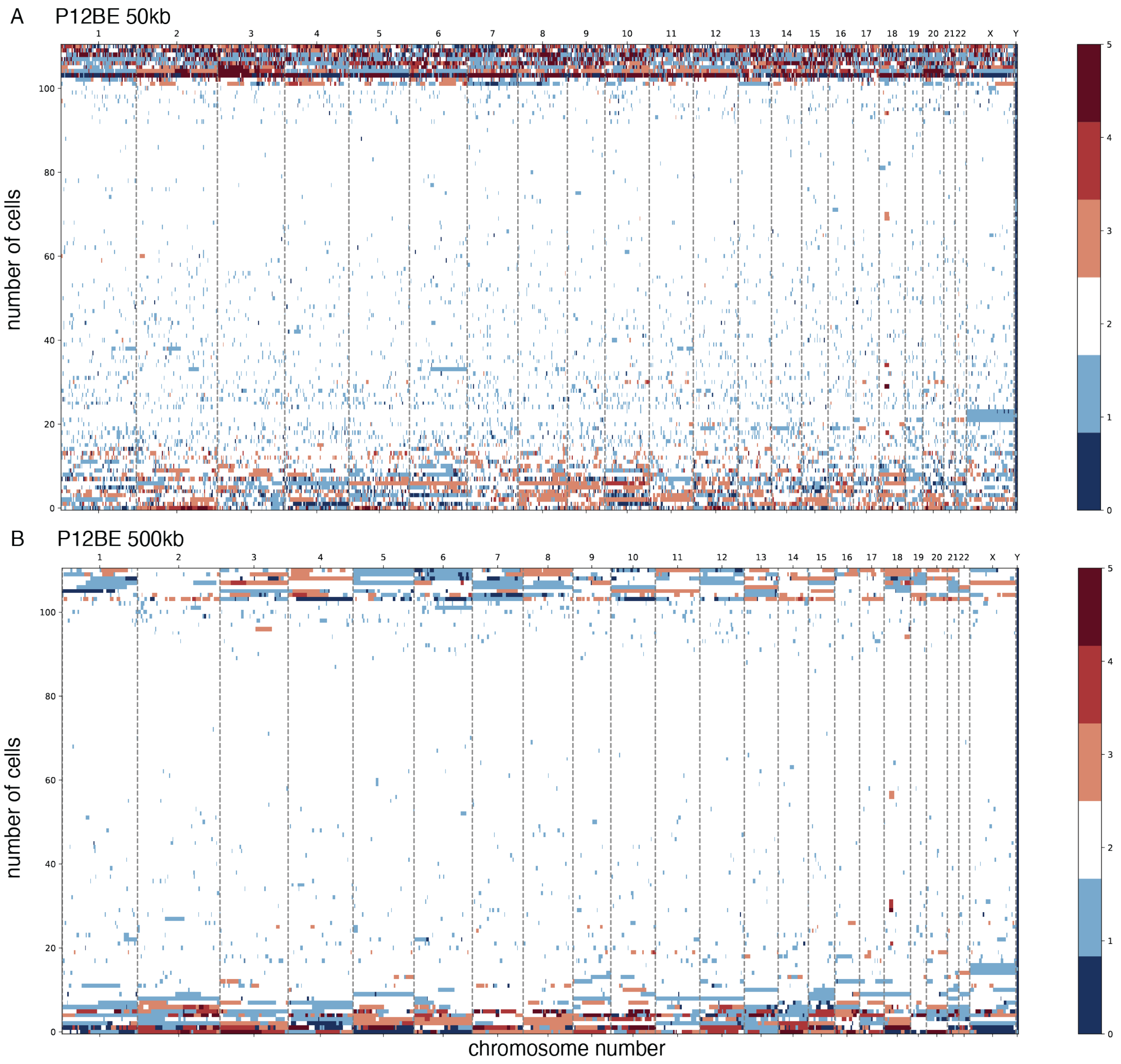


**Fig. S10. Single nucleus Copy Number Heatmap for P12.** BE: Barrett's esophagus. X axis: genome coordinates, upper: chromosome number. Y axis: cell number. Color scale: copy numbers. White=2, Orange=3, red=4, dark red=5, lighter blue=1, dark blue=0. **A: 50kb bin; B: 500kb bin.**

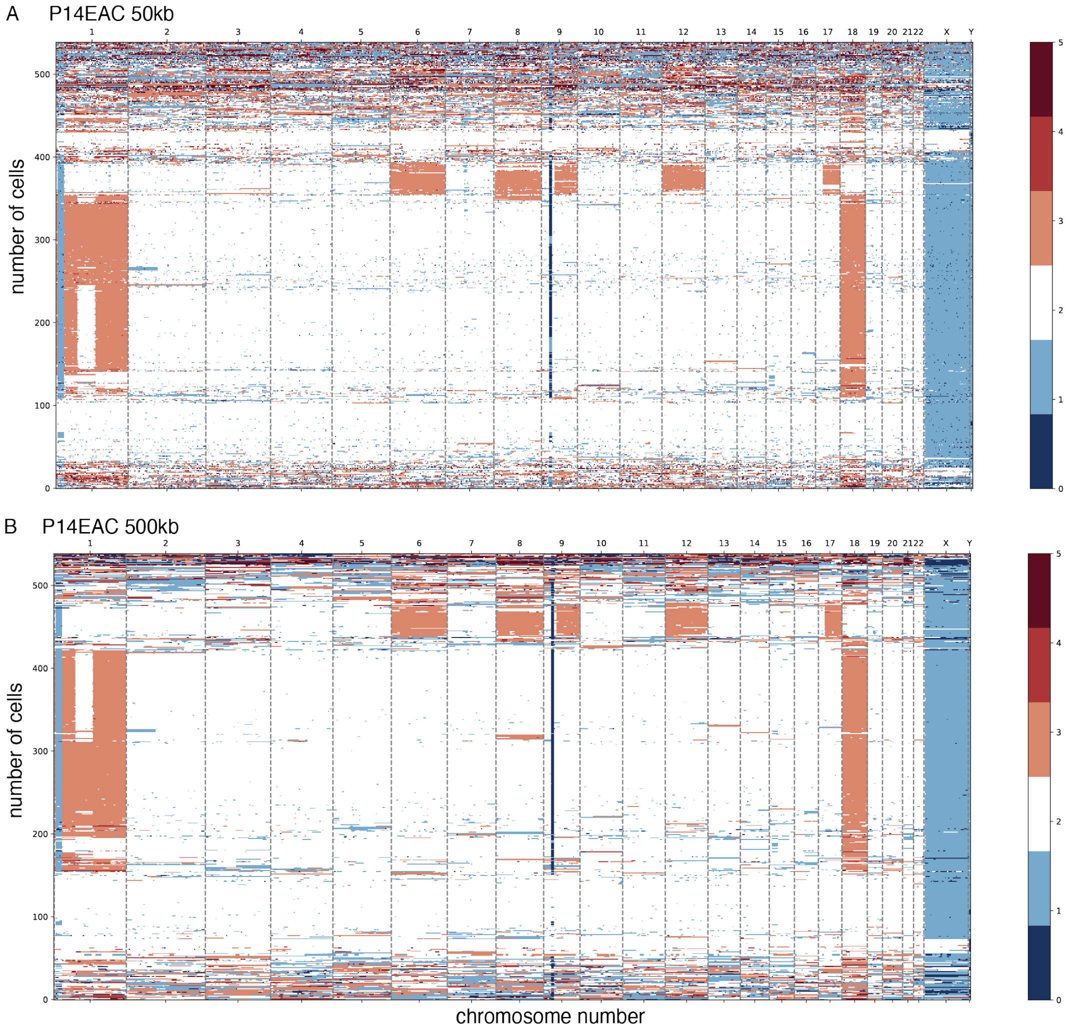


**Fig. S11. Single nucleus Copy Number Heatmap for P14.** EAC: esophageal adenocarcinoma, X axis: genome coordinates, upper: chromosome number. Y axis: cell number. Color scale: copy numbers. White=2, Orange=3, red=4, dark red=5, lighter blue=1, dark blue=0. **A: 50kb bin; B: 500kb bin.**

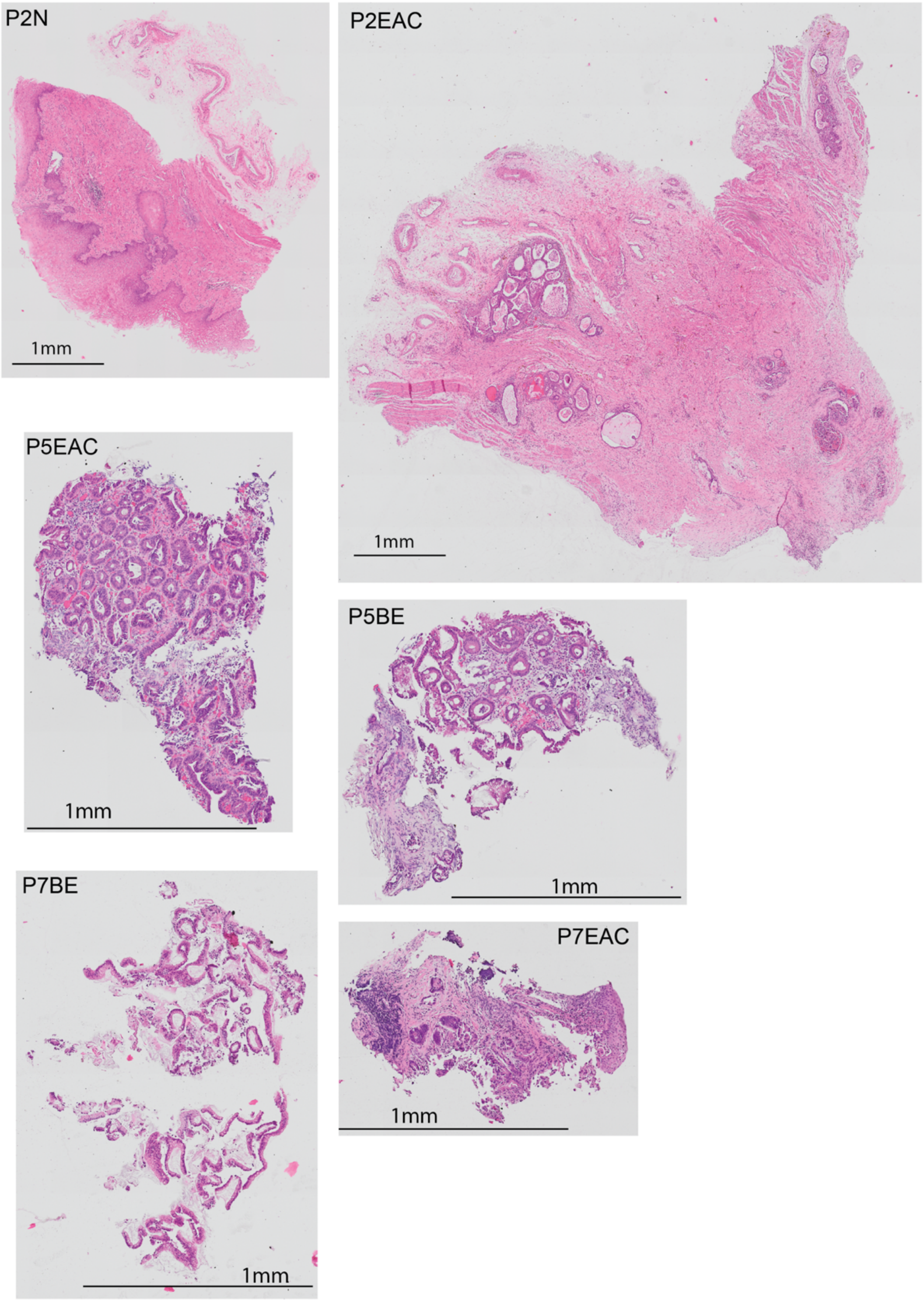


**Fig. S12. Pathological impressions of selected tissue slides used in this study.** P2EAC: Foci of invasive adenocarcinoma in the submucosal tissue of the esophagus. P2N: Normal stratified squamous esophageal mucosa. P5EAC: Barrett's esophagus with low-grade dysplasia in a patient with a clinical history of adenocarcinoma. P5BE: Barrett's esophagus with focal dysplasia. P7EAC: Invasive adenocarcinoma underlying esophageal squamous mucosa. P7BE: Barrett's esophagus without dysplasia. Scale bars: 1mm.

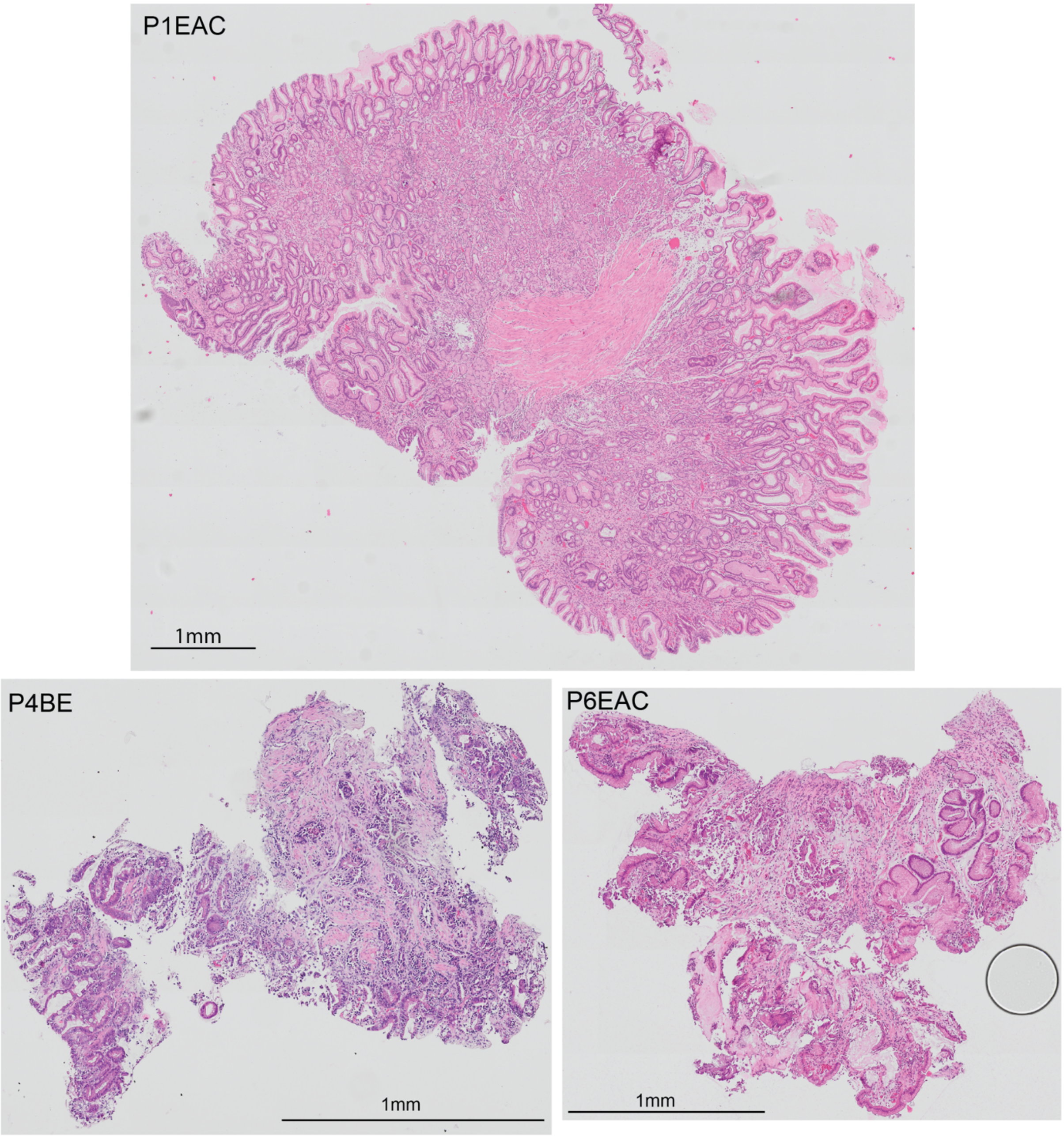


**Fig. S13. Pathological impressions of selected tissue slides used in this study (continued).** P1EAC: Tumor site following neoadjuvant therapy showing gastric type glandular mucosa with focal areas of low-grade dysplasia. No evidence of residual invasive adenocarcinoma. P4BE: Barrett's esophagus with high-grade dysplasia and area of suspicious for invasive adenocarcinoma. P6EAC: Invasive adenocarcinoma underlying gastric type mucosa. Scale bars: 1mm.

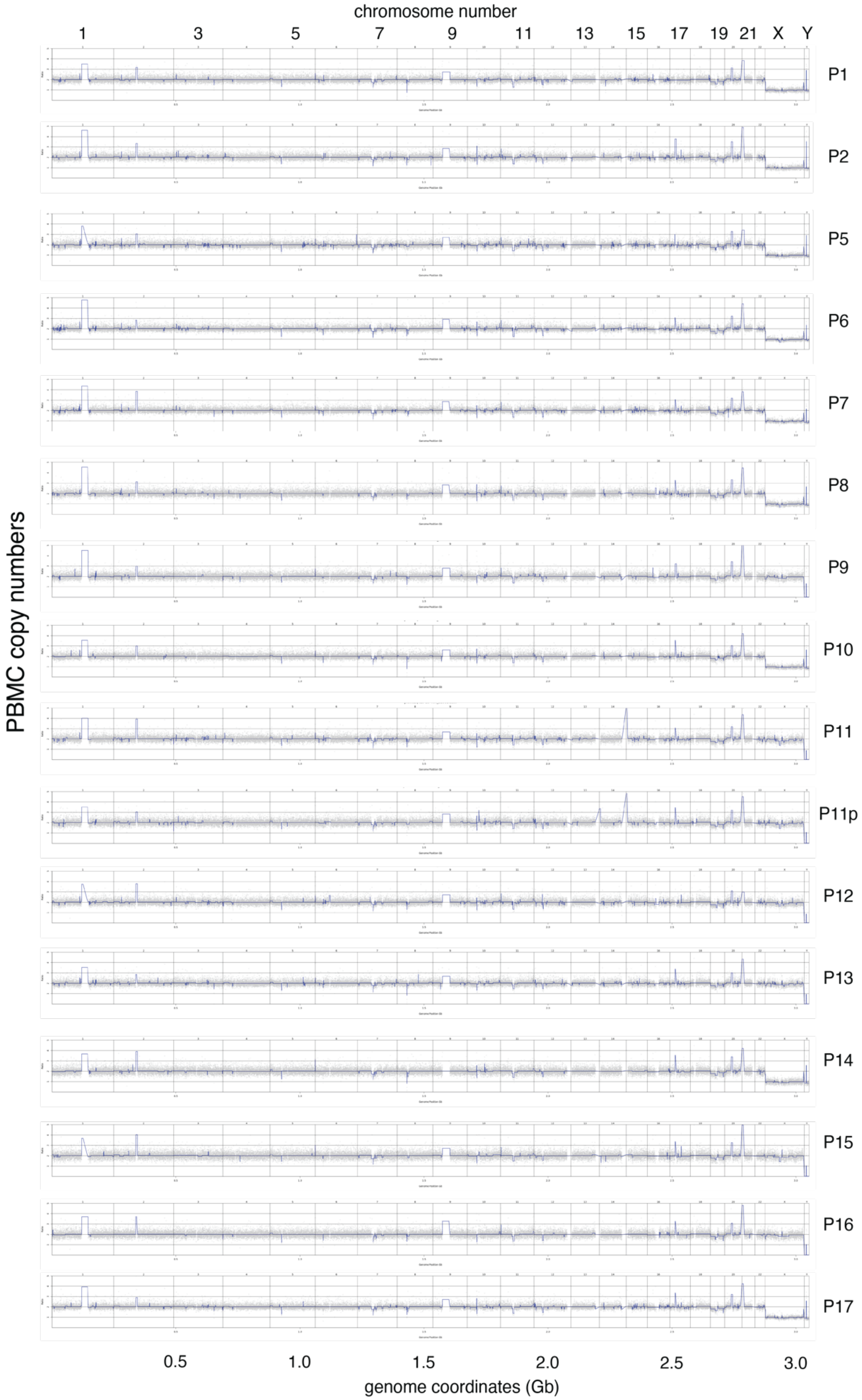


**Fig. S14. Y chromosome loss does not originate from the germline.** Our findings suggest that Y chromosome loss (Y loss) is not a germline event but rather a somatic alteration. To investigate this, we performed whole-genome sequencing on genomic DNA (gDNA) extracted from peripheral blood mononuclear cells (PBMCs). The analysis revealed a distinct lack of major copy number alterations in these PBMC-derived genomes, contrasting sharply with the more complex genomic landscapes observed in tumor samples and tumor-adjacent normal tissues. The relative genomic stability of the PBMC-extracted gDNA, which we describe as "quieter," further supports the hypothesis that Y loss occurs as a post-zygotic event. This somatic origin of Y loss aligns with its observed prevalence in certain subpopulations and its association with other somatic mutations and alterations within the tumor microenvironment.

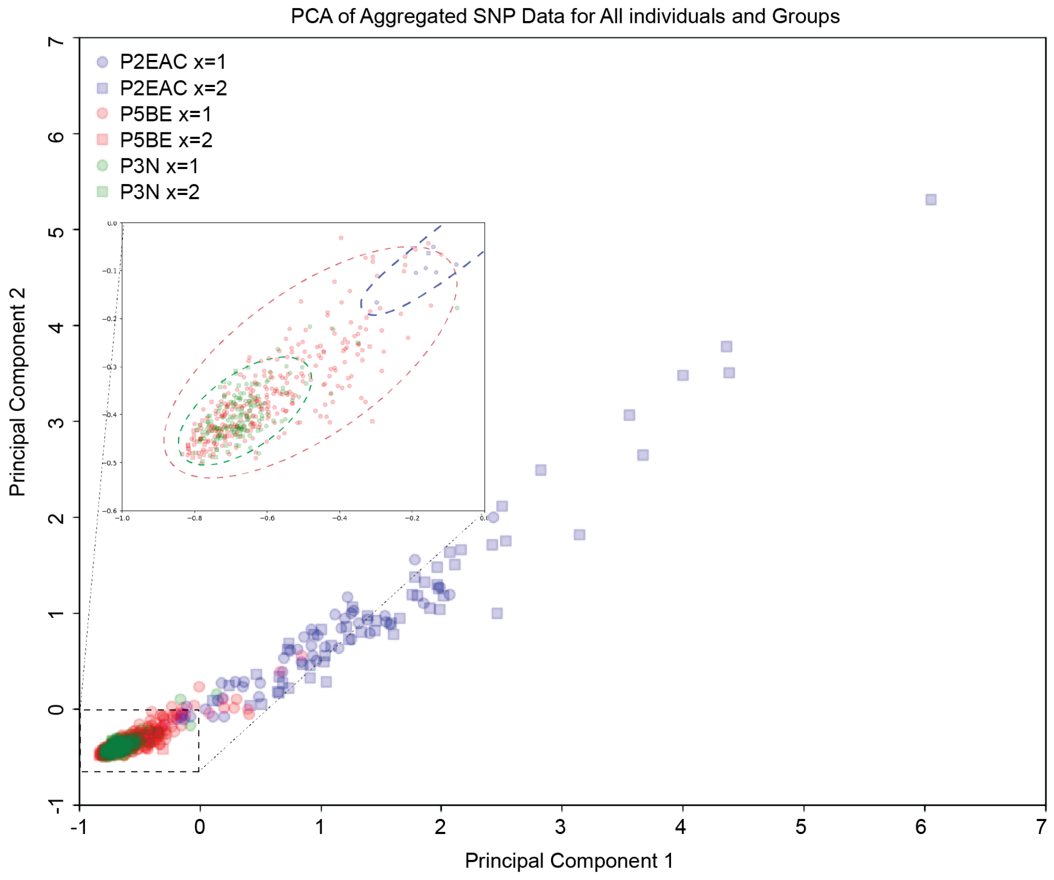


**Fig. S15. PCA analysis of SNP from specimen with X duplications.** PCA analysis suggested that the X duplicated cells and non-duplicated cells were most likely to come from the same individual and not due to contaminations. Circle: x chromosome copy number =1; Square: x chromosome copy number =2.

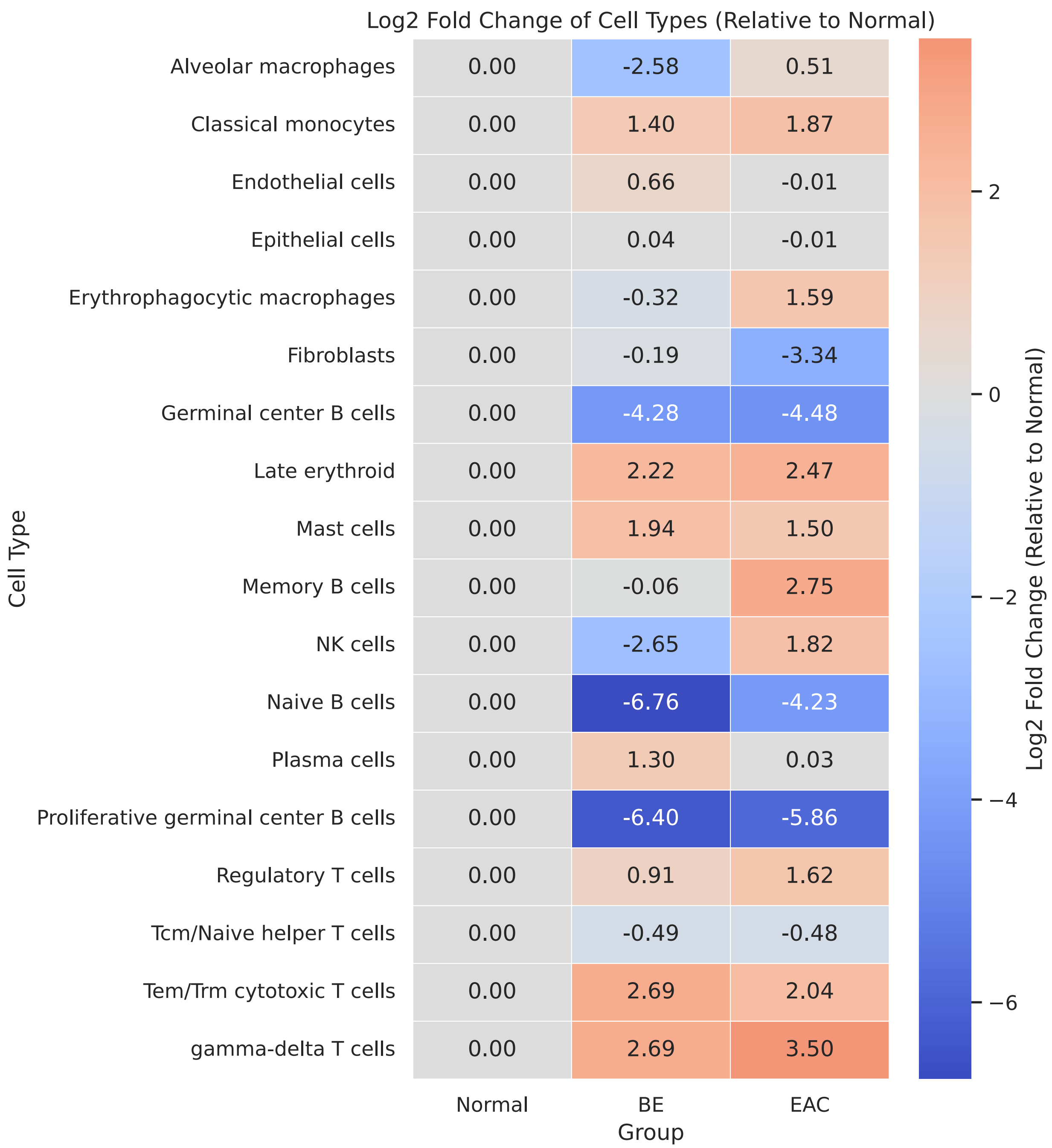


**Fig. S16. Heatmap analysis of cell compositions in EAC/BE and normal groups.** Red: increased proportion compared to normal. Blue: decreased proportion.

5

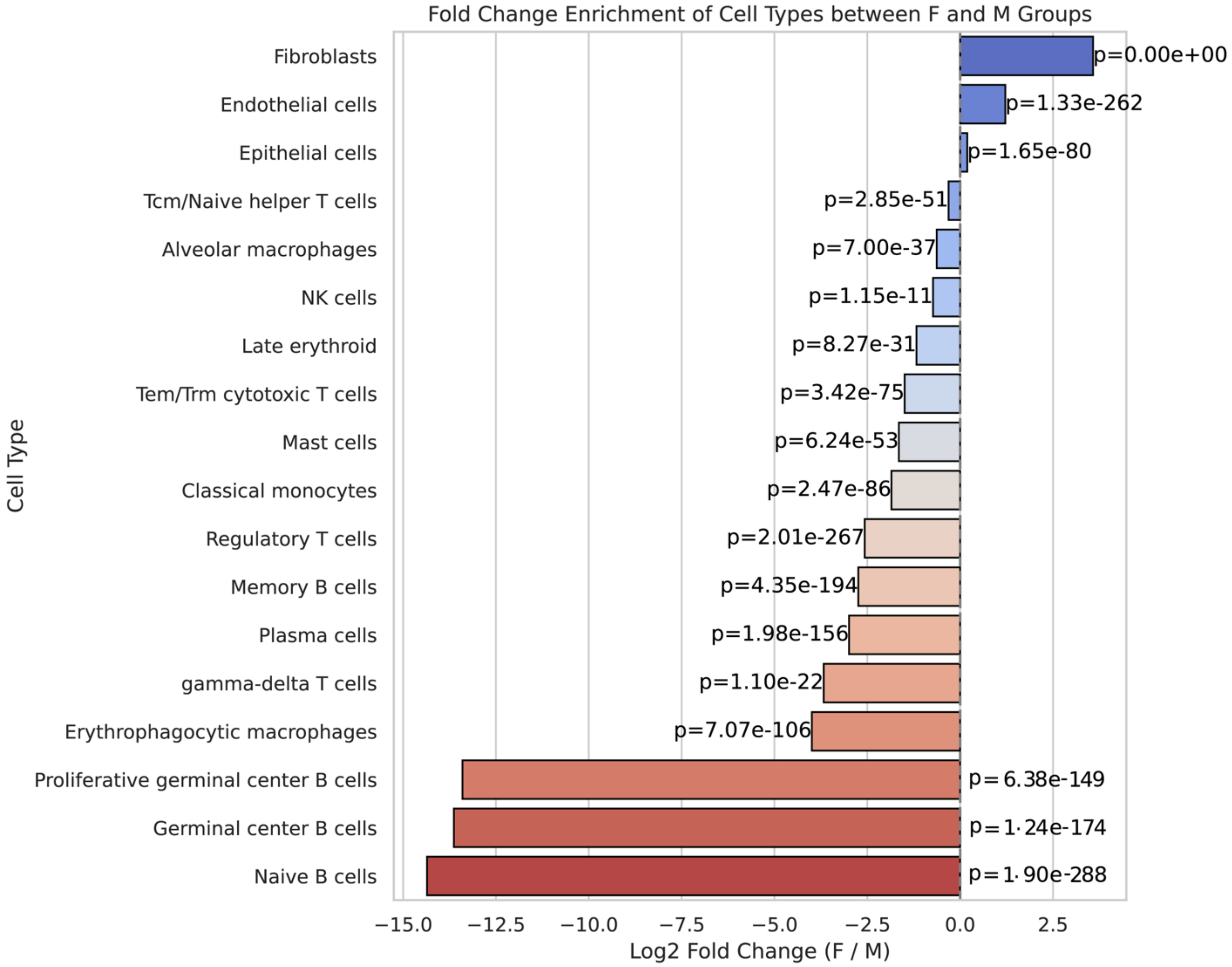


**Fig. S17. cell compositions in Females vs male groups.** Red: cell types depleted in females. Blue: cell types enriched in females.

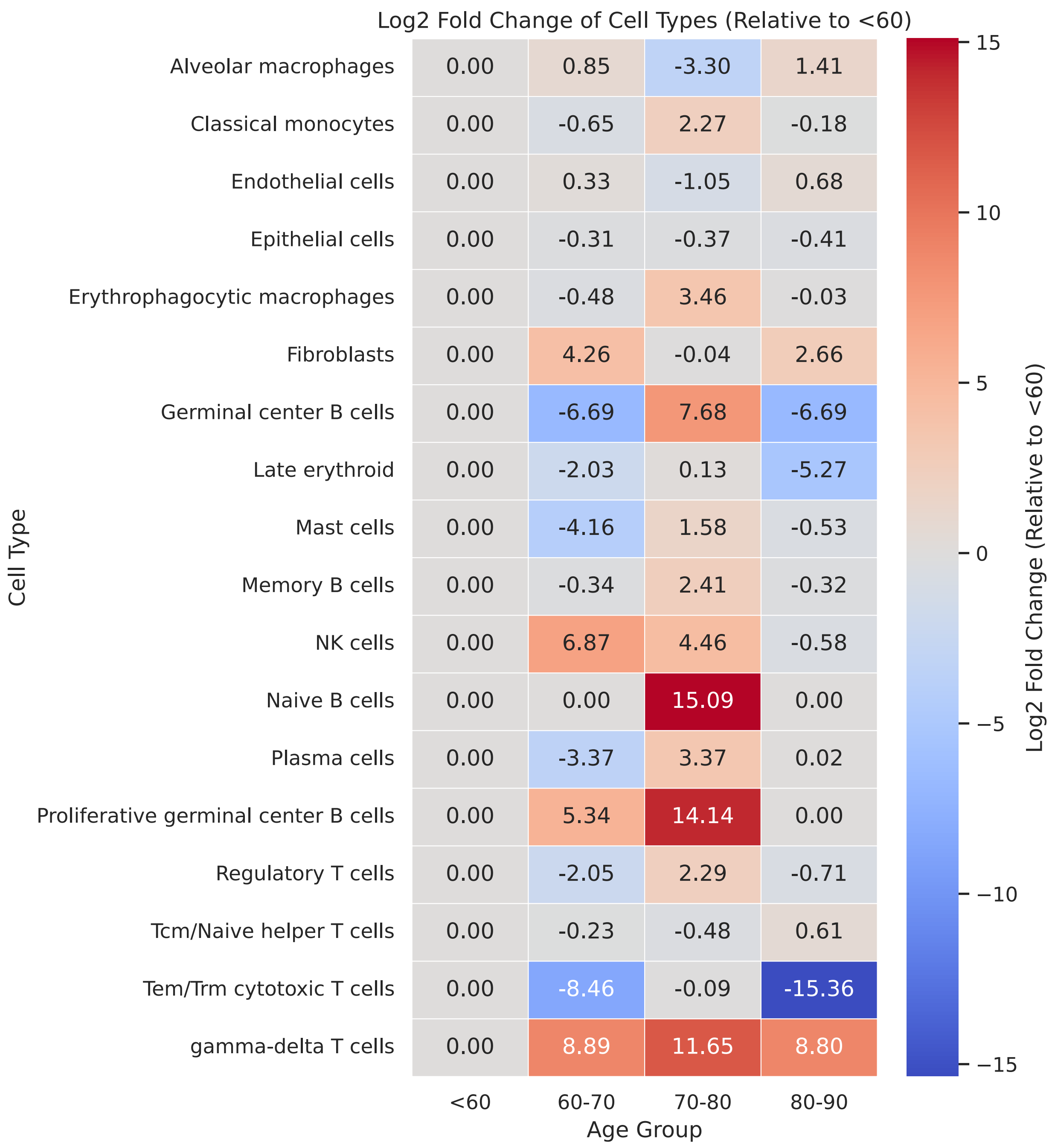


**Fig. S18. Heatmap analysis of cell compositions in different age groups.** Red: increased proportion compared to <60. Blue: decreased proportion compared to <60 group.

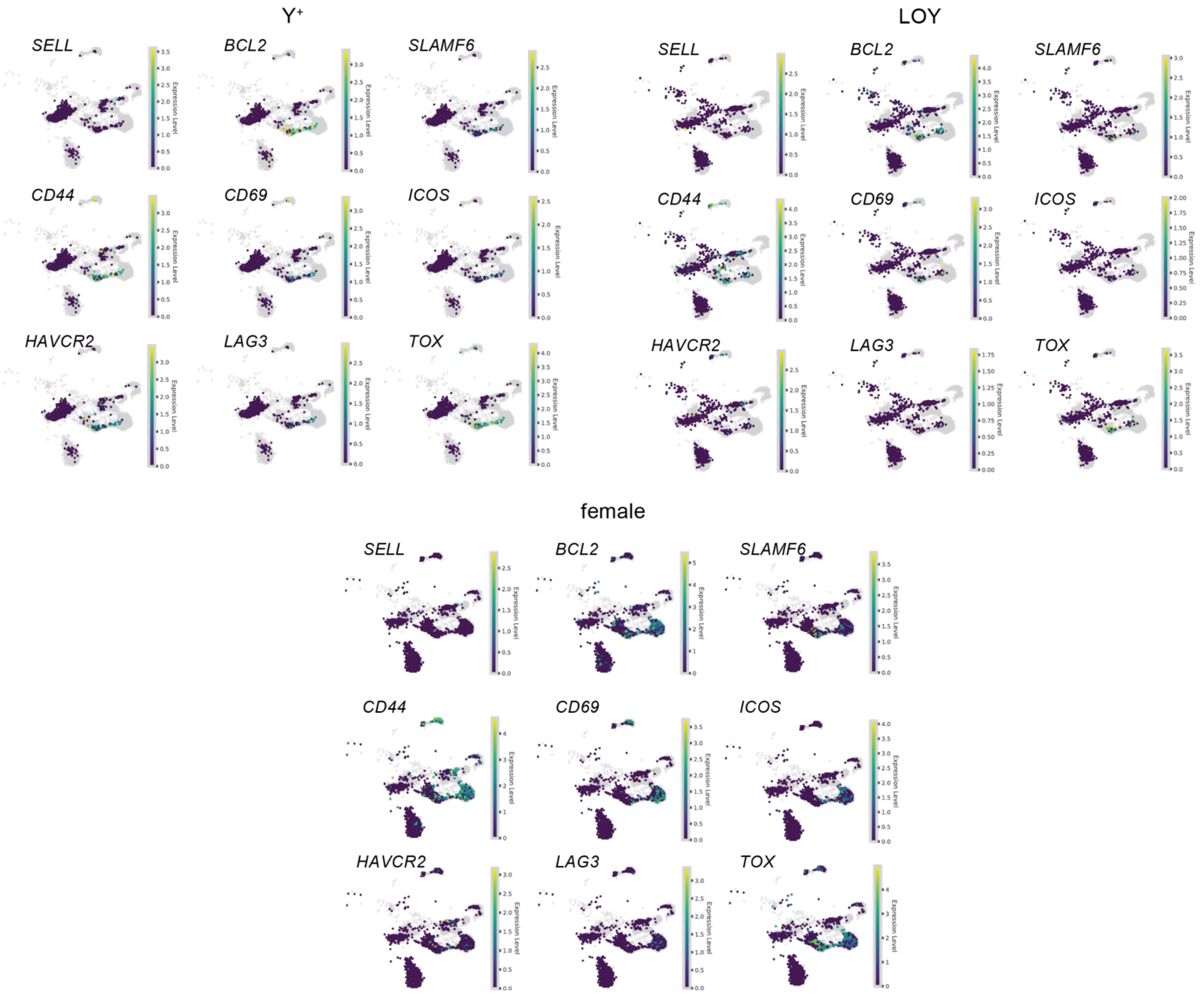


**Fig. S19. T cell exhaustion markers were not increased in Loss of Y groups.** T cell stem-like (*SELL*, *BCL2*, *SLAMF6*), activation (*CD44*, *CD69*, *ICOS*), and exhaustion (*TIM3*, *LAG3*, *TOX*) markers were compared between Y+, LOY, and female groups.

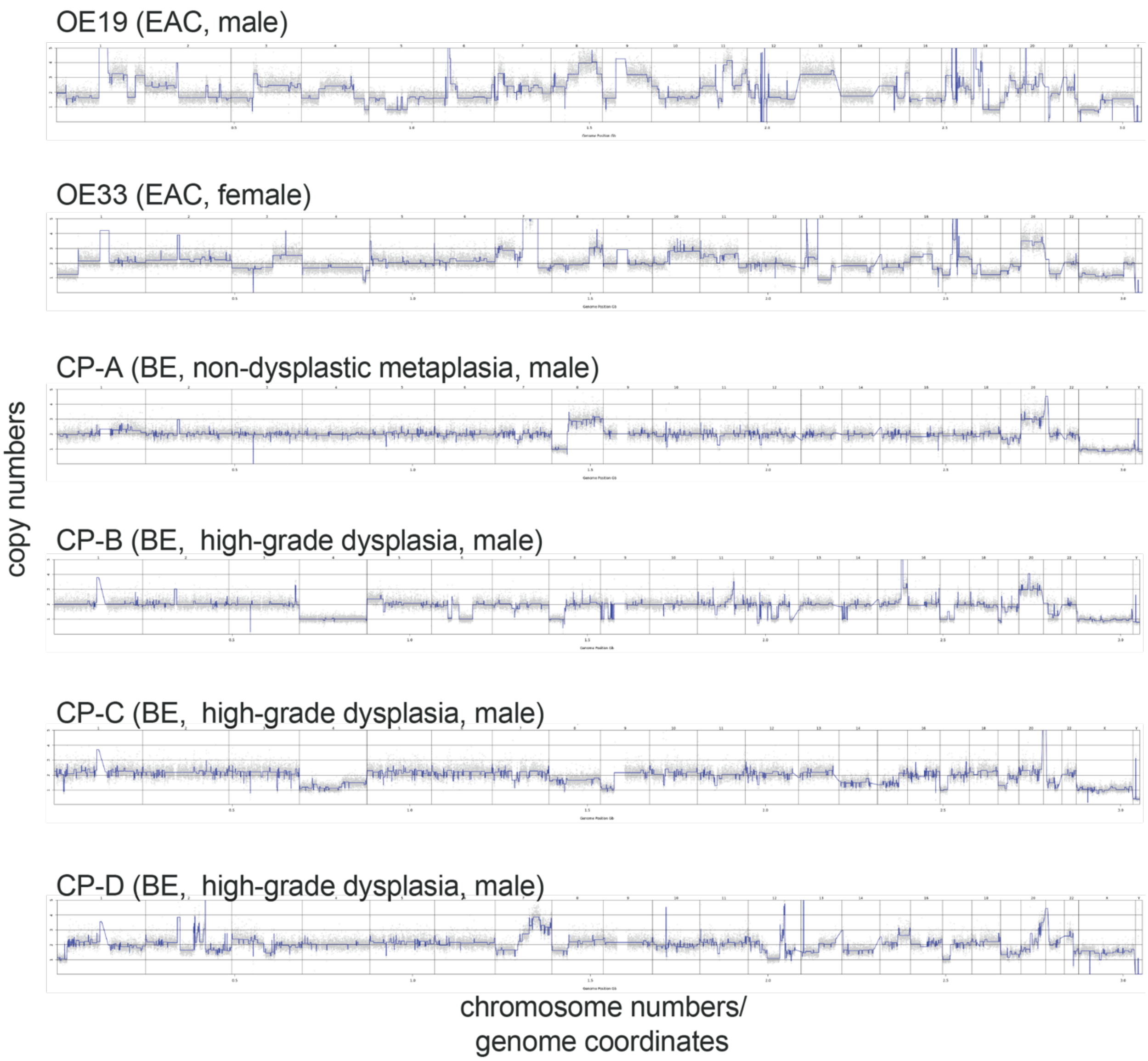


**Fig. 20. Shallow whole-genome sequencing (WGS) copy number analysis of common Barrett's esophagus (BE) and esophageal adenocarcinoma (EAC) cell lines.** CP-A, CP-B, CP-C, and CP-D are male-derived BE cell lines. CP-A retains chromosome Y, whereas CP-B and CP-C show partial Y loss. CP-D displays complete Y loss and elevated X dosage, indicating that a portion of the cells is likely duplicated. OE19, a male derived EAC cell line, has complete loss of Y (LOY) and elevated dosage of an arm of the X chromosome. OE33, a female derived EAC cell line, serves as a control.

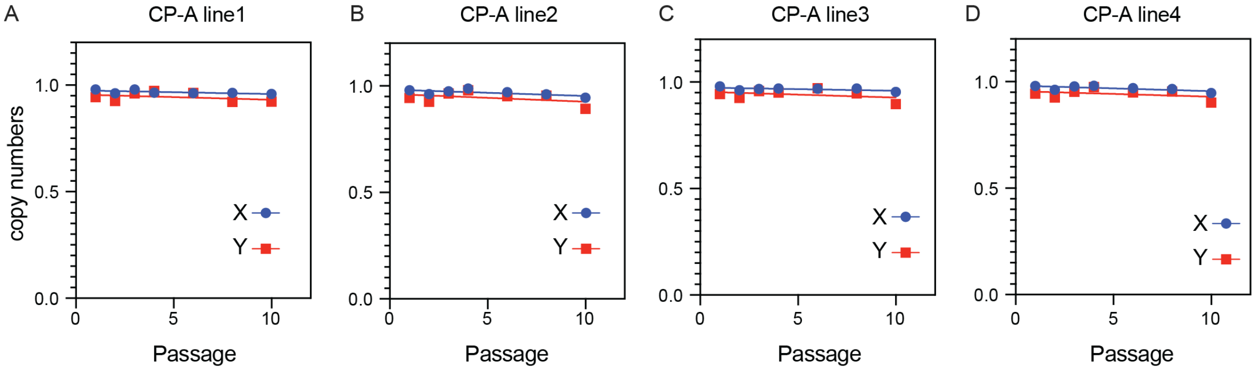


**Fig. S21.** Shallow whole-genome sequencing (WGS) copy-number analysis of the male-derived Barrett’s esophagus (BE) cell line CP-A shows stable X and Y chromosome dosage over time.

**Table S1. Data Availability of Single-Cell, Bulk Genomic and FISH Assays, Along with Clinical Characteristics of the study**

Table S1. Data Availability of Single-Cell, Bulk Genomic and FISH Assays, Along with Clinical Characteristics of the study

| patient number | UltraCNA | 10x | Blood_shallowWGS | FISH | obesity status | age group | Sex |
|---|---|---|---|---|---|---|---|
| P1 | + (EAC) | - | + | | not obese | 70-80 | M |
| P2 | +(EAC,N) | +(EAC, N) | + | +(EAC,N) | not obese | 60-70 | M |
| P3 | +(EAC,N) | +(EAC, N) | - | - | obesity | <60 | M |
| P4 | +(BE,N) | +(N) | - | - | not obese | 60-70 | M |
| P5 | +(EAC,BE, N) | +(EAC,BE, N) | + | +(BE,N) | not obese | 80-90 | M |
| P6 | +(EAC,N) | +(EAC,N) | + | - | not obese | 70-80 | M |
| P7 | +(EAC, BE, N) | - | + | +(EAC,BE) | not obese | 60-70 | M |
| P8 | +(EAC, BE) | +(EAC,BE, N) | + | +(EAC,BE, N) | obesity | 60-70 | M |
| P9 | - | +(BE, N) | + | - | obesity | <60 | F |
| P10 | - | +(BE, N) | + | - | obesity | 80-90 | F |
| P11 | +(EAC) | +(EAC) | + | +(EAC) | obesity | 60-70 | F |
| P11p | - | +(EAC,N) | + | - | obesity | 60-70 | F |
| P12 | +(BE) | - | + | - | not obese | 60-70 | F |
| P13 | - | +(EAC) | + | - | not obese | 60-70 | F |
| P14 | +(EAC) | +(EAC*2, N) | + | - | not obese | 80-90 | M |
| P15 | - | +(BE) | + | - | not obese | 70-80 | F |
| P16 | - | +(BE) | + | - | not obese | <60 | F |
| P17 | - | +(EAC, BE, N) | + | - | not obese | 70-80 | M |
| P17p | - | +(EAC, BE, N) | - | - | not obese | 70-80 | M |

In this table, the presence of 10x Genomics or ultraCNV information simply indicates that these analyses were performed for those specific samples. The absence of data for a sample does not mean the specimen itself was unavailable; rather, it reflects that we did not generate those particular analyses for that specimen. The table is intended only to denote where these specific data types exist, not to indicate the presence or absence of the underlying specimens. Specimen that originally have at least 250 nucleus were kept. Specimens with signs of contamination and/or very low quality were discarded.

**Table S2. Single-Cell RNA-Seq Sample Grouping Information.**

| Table S2. Single-Cell RNA-Seq Sample Grouping Information. | | | | | | | |
|---|---|---|---|---|---|---|---|
| annotation | group1 | group2 | group3 | group4 | group5 | group6 | group7 |
| P2EAC | EAC | P2 | v3 | M | LOY | not obese | 60-70 |
| P2N | Normal | P2 | v3 | M | LOY | not obese | 60-70 |
| P3EAC | EAC | P3 | v3 | M | Y+ | obese | <60 |
| P3N | Normal | P3 | v3 | M | LOY | obese | <60 |
| P4N | Normal | P4 | v3 | M | Y+ | not obese | 60-70 |
| P5BE | BE | P5 | v4 | M | LOY | not obese | 80-90 |
| P5EAC | EAC | P5 | v3 | M | LOY | not obese | 80-90 |
| P5N | Normal | P5 | v3 | M | LOY | not obese | 80-90 |
| P6EAC | EAC | P6 | v3 | M | Y+ | not obese | 70-80 |
| P6N | Normal | P6 | v3 | M | Y+ | not obese | 70-80 |
| P8BE | BE | P8 | v3 | M | Y+ | obese | 60-70 |
| P8EAC | EAC | P8 | v3 | M | LOY | obese | 60-70 |
| P8N | Normal | P8 | v3 | M | not determined | obese | 60-70 |
| P9BE | BE | P9 | v4 | F | Female | obese | <60 |
| P9N | Normal | P9 | v4 | F | Female | obese | <60 |
| P10BE | BE | P10 | v4 | F | Female | obese | 80-90 |
| P10N | Normal | P10 | v4 | F | Female | obese | 80-90 |
| P11EAC | EAC | P11 | v3 | F | Female | obese | 60-70 |
| P11pEAC | EAC | P11 | v3 | F | Female | obese | 60-70 |
| P11pN | Normal | P11 | v3 | F | Female | obese | 60-70 |
| P13EAC | EAC | P13 | v3 | F | Female | not obese | 60-70 |
| P14EAC#1 | EAC | P14 | v3 | M | Y+ | not obese | 80-90 |
| P14EAC#2 | EAC | P14 | v3 | M | Y+ | not obese | 80-90 |
| P14N | Normal | P14 | v3 | M | not determined | not obese | 80-90 |
| P15BE | BE | P15 | v4 | F | Female | not obese | 70-80 |
| P16BE | BE | P16 | v3 | F | Female | not obese | <60 |
| P17BE | BE | P17 | v4 | M | not determined | not obese | 70-80 |
| P17N | Normal | P17 | v4 | M | not determined | not obese | 70-80 |
| P17EAC | EAC | P17 | v4 | M | not determined | not obese | 70-80 |

| | | | | | | | |
|---|---|---|---|---|---|---|---|
| P17pBE | BE | P17 | v4 | M | not determined | not obese | 70-80 |
| P17pEAC | EAC | P17 | v4 | M | not determined | not obese | 70-80 |
| P17pN | Normal | P17 | v4 | M | not determined | not obese | 70-80 |
| Groups 1–7 indicate (1) sample type, where EAC represents esophageal adenocarcinoma, Normal denotes normal esophageal mucosa, and BE indicates Barrett's esophagus; (2) patient ID, specifying from which patient the sample was obtained; (3) 10x 3′ single-cell RNA-Seq version, indicating which version of the 10x Genomics 3′ RNA-seq platform was used; (4) sex, where M is male and F is female; (5) Y status, based on single-cell Ultra-CNA data (e.g., presence or absence of the loss of Y chromosome, LOY: more than 15% of single-nuclei lost >80% of Y chromosome using either 50kb or 500kb bin); (6) obesity status, categorized as obese for BMI > 30; and (7) age group. | | | | | | | |

**Table S3. Primer information**

| Sequence Name | Sequence |
|---|---|
| N703 | CAA GCA GAA GAC GGC ATA CGA GAT TTC TGC CTG TCT CGT GGG CTC GG |
| N704 | CAA GCA GAA GAC GGC ATA CGA GAT GCT CAG GAG TCT CGT GGG CTC GG |
| N705 | CAA GCA GAA GAC GGC ATA CGA GAT AGG AGT CCG TCT CGT GGG CTC GG |
| N706 | CAA GCA GAA GAC GGC ATA CGA GAT CAT GCC TAG TCT CGT GGG CTC GG |
| N707 | CAA GCA GAA GAC GGC ATA CGA GAT GTA GAG AGG TCT CGT GGG CTC GG |
| N708 | CAA GCA GAA GAC GGC ATA CGA GAT CCT CTC TGG TCT CGT GGG CTC GG |
| N709 | CAA GCA GAA GAC GGC ATA CGA GAT AGC GTA GCG TCT CGT GGG CTC GG |
| N710 | CAA GCA GAA GAC GGC ATA CGA GAT CAG CCT CGG TCT CGT GGG CTC GG |
| N711 | CAA GCA GAA GAC GGC ATA CGA GAT TGC CTC TTG TCT CGT GGG CTC GG |
| N712 | CAA GCA GAA GAC GGC ATA CGA GAT TCC TCT ACG TCT CGT GGG CTC GG |
| N714 | CAA GCA GAA GAC GGC ATA CGA GAT TCA TGA GCG TCT CGT GGG CTC GG |
| N715 | CAA GCA GAA GAC GGC ATA CGA GAT CCT GAG ATG TCT CGT GGG CTC GG |
| N716 | CAA GCA GAA GAC GGC ATA CGA GAT TAG CGA GTG TCT CGT GGG CTC GG |
| N718 | CAA GCA GAA GAC GGC ATA CGA GAT GTA GCT CCG TCT CGT GGG CTC GG |
| N719 | CAA GCA GAA GAC GGC ATA CGA GAT TAC TAC GCG TCT CGT GGG CTC GG |
| N720 | CAA GCA GAA GAC GGC ATA CGA GAT AGG CTC CGG TCT CGT GGG CTC GG |
| N722 | CAA GCA GAA GAC GGC ATA CGA GAT CTG CGC ATG TCT CGT GGG CTC GG |
| N723 | CAA GCA GAA GAC GGC ATA CGA GAT GAG CGC TAG TCT CGT GGG CTC GG |

| | |
|---|---|
| N724 | CAA GCA GAA GAC GGC ATA CGA GAT CGC TCA GTG TCT CGT GGG CTC GG |
| N726 | CAA GCA GAA GAC GGC ATA CGA GAT GTC TTA GGG TCT CGT GGG CTC GG |
| N727 | CAA GCA GAA GAC GGC ATA CGA GAT ACT GAT CGG TCT CGT GGG CTC GG |
| N728 | CAA GCA GAA GAC GGC ATA CGA GAT TAG CTG CAG TCT CGT GGG CTC GG |
| N731 | CAA GCA GAA GAC GGC ATA CGA GAT ATC ACG TTG TCT CGT GGG CTC GG |
| N732 | CAA GCA GAA GAC GGC ATA CGA GAT CGA TGT TTG TCT CGT GGG CTC GG |
| N733 | CAA GCA GAA GAC GGC ATA CGA GAT TTA GGC ATG TCT CGT GGG CTC GG |
| N734 | CAA GCA GAA GAC GGC ATA CGA GAT TGA CCA CTG TCT CGT GGG CTC GG |
| N735 | CAA GCA GAA GAC GGC ATA CGA GAT ACA GTG GTG TCT CGT GGG CTC GG |
| N736 | CAA GCA GAA GAC GGC ATA CGA GAT GCC AAT GTG TCT CGT GGG CTC GG |
| N737 | CAA GCA GAA GAC GGC ATA CGA GAT CAG ATC TGG TCT CGT GGG CTC GG |
| N738 | CAA GCA GAA GAC GGC ATA CGA GAT ACT TGA TGG TCT CGT GGG CTC GG |
| N739 | CAA GCA GAA GAC GGC ATA CGA GAT GAT CAG CGG TCT CGT GGG CTC GG |
| N740 | CAA GCA GAA GAC GGC ATA CGA GAT TAG CTT GTG TCT CGT GGG CTC GG |
| S504 | AAT GAT ACG GCG ACC ACC GAG ATC TAC ACA GAG TAG ATC GTC GGC AGC GTC |
| S505 | AAT GAT ACG GCG ACC ACC GAG ATC TAC ACG TAA GGA GTC GTC GGC AGC GTC |
| S506 | AAT GAT ACG GCG ACC ACC GAG ATC TAC ACA CTG CAT ATC GTC GGC AGC GTC |
| S507 | AAT GAT ACG GCG ACC ACC GAG ATC TAC ACA AGG AGT ATC GTC GGC AGC GTC |
| S508 | AAT GAT ACG GCG ACC ACC GAG ATC TAC ACC TAA GCC TTC GTC GGC AGC GTC |
| S510 | AAT GAT ACG GCG ACC ACC GAG ATC TAC ACC GTC TAA TTC GTC GGC AGC GTC |
| S511 | AAT GAT ACG GCG ACC ACC GAG ATC TAC ACT CTC TCC GTC GTC GGC AGC GTC |
| S513 | AAT GAT ACG GCG ACC ACC GAG ATC TAC ACT CGA CTA GTC GTC GGC AGC GTC |
| S515 | AAT GAT ACG GCG ACC ACC GAG ATC TAC ACT TCT AGC TTC GTC GGC AGC GTC |
| S516 | AAT GAT ACG GCG ACC ACC GAG ATC TAC ACC CTA GAG TTC GTC GGC AGC GTC |
| S517 | AAT GAT ACG GCG ACC ACC GAG ATC TAC ACG CGT AAG ATC GTC GGC AGC GTC |
| S518 | AAT GAT ACG GCG ACC ACC GAG ATC TAC ACC TAT TAA GTC GTC GGC AGC GTC |

| | |
|---|---|
| S520 | AAT GAT ACG GCG ACC ACC GAG ATC TAC ACA AGG CTA TTC GTC GGC AGC GTC |
| S521 | AAT GAT ACG GCG ACC ACC GAG ATC TAC ACG AGC CTT ATC GTC GGC AGC GTC |
| S522 | AAT GAT ACG GCG ACC ACC GAG ATC TAC ACT TAT GCG ATC GTC GGC AGC GTC |
| S525 | AAT GAT ACG GCG ACC ACC GAG ATC TAC ACT AAG CGT TTC GTC GGC AGC GTC |
| S526 | AAT GAT ACG GCG ACC ACC GAG ATC TAC ACT CCG TCT TTC GTC GGC AGC GTC |
| S527 | AAT GAT ACG GCG ACC ACC GAG ATC TAC ACT GTA CCT TTC GTC GGC AGC GTC |
| S528 | AAT GAT ACG GCG ACC ACC GAG ATC TAC ACT TCT GTG TTC GTC GGC AGC GTC |
| S529 | AAT GAT ACG GCG ACC ACC GAG ATC TAC ACT CTG CTG TTC GTC GGC AGC GTC |
| S530 | AAT GAT ACG GCG ACC ACC GAG ATC TAC ACT TGG AGG TTC GTC GGC AGC GTC |
| S531 | AAT GAT ACG GCG ACC ACC GAG ATC TAC ACT CGA GCG TTC GTC GGC AGC GTC |
| S532 | AAT GAT ACG GCG ACC ACC GAG ATC TAC ACT GAT ACG TTC GTC GGC AGC GTC |
| S533 | AAT GAT ACG GCG ACC ACC GAG ATC TAC ACT GCA TAG TTC GTC GGC AGC GTC |
| S534 | AAT GAT ACG GCG ACC ACC GAG ATC TAC ACT TGA CTC TTC GTC GGC AGC GTC |
| S535 | AAT GAT ACG GCG ACC ACC GAG ATC TAC ACT GCG ATC TTC GTC GGC AGC GTC |
| S536 | AAT GAT ACG GCG ACC ACC GAG ATC TAC ACT TCC TGC TTC GTC GGC AGC GTC |
| S537 | AAT GAT ACG GCG ACC ACC GAG ATC TAC ACT AGT GAC TTC GTC GGC AGC GTC |
| S538 | AAT GAT ACG GCG ACC ACC GAG ATC TAC ACT ACA GGA TTC GTC GGC AGC GTC |
| S539 | AAT GAT ACG GCG ACC ACC GAG ATC TAC ACT CCT CAA TTC GTC GGC AGC GTC |
| S540 | AAT GAT ACG GCG ACC ACC GAG ATC TAC ACT GTG GTT GTC GTC GGC AGC GTC |
| S541 | AAT GAT ACG GCG ACC ACC GAG ATC TAC ACT ACT AGT CTC GTC GGC AGC GTC |
| S542 | AAT GAT ACG GCG ACC ACC GAG ATC TAC ACT TCC ATT GTC GTC GGC AGC GTC |
| S543 | AAT GAT ACG GCG ACC ACC GAG ATC TAC ACT CGA AGT GTC GTC GGC AGC GTC |
| S544 | AAT GAT ACG GCG ACC ACC GAG ATC TAC ACT AAC GCT GTC GTC GGC AGC GTC |

| | |
|---|---|
| S545 | AAT GAT ACG GCG ACC ACC GAG ATC TAC ACT TGG TAT GTC GTC GGC AGC GTC |
| S546 | AAT GAT ACG GCG ACC ACC GAG ATC TAC ACT GAA CTG GTC GTC GGC AGC GTC |
| S547 | AAT GAT ACG GCG ACC ACC GAG ATC TAC ACT ACT TCG GTC GTC GGC AGC GTC |
| S548 | AAT GAT ACG GCG ACC ACC GAG ATC TAC ACT CTC ACG GTC GTC GGC AGC GTC |
